# The tetrapod fauna of the upper Permian Naobaogou Formation of China: a new mid-sized pareiasaur *Yinshanosaurus angustus* gen. et sp. nov. and its implication for the phylogenetic relationship of pareiasaurs

**DOI:** 10.1101/2024.11.23.624968

**Authors:** Jian Yi, Jun Liu

## Abstract

Pareiasauria is a specialized clade of herbivorous tetrapods that existed throughout Pangaea during the mid to late Permian period. The phylogenetic relationships of Chinese pareiasaur species remain controversy for several decades, primarily due to the poor preservation of known specimens. Until the report of *Shihtienfenia completus* in 2019, no complete skull had been documented for Chinese pareiasaurs. The present study describes a mid-sized pareiasaur, *Yinshanosaurus angustus* gen. et sp. nov., based on a nearly complete skull and an articulated partial postcranial skeleton collected from the Naobaogou Formation in 2018. It presents several significant new morphological features such as high and narrow skull, with the length of the skull table exceeding the width between the two quadratojugals; snout dimension as high as wide; long frontal with a length-to-width ratio of ∼3.0; U-shaped paraoccipital process in occipital view; and maxillary teeth oriented vertically, with only 7-9 cusps. Although the phylogenetic framework of pareiasaurs still requires further refinement, the current analysis yields distinct phylogenetic positions for Chinese pareiasaurs and establishes a new monophyletic clade that includes *S*. *completus* and *Y*. *angustus*.

---

PAREIASAURIA is a bizarre herbivorous clade of tetrapods that exist in the Guadalupian and Lopingian (Smith *et al*. 2020) and was a victim at both the late Capitanian (Day *et al*. 2015) and the end-Permian mass extinction events (Smith and Botha-Brink 2014). Pareiasauria exhibits worldwide distribution, with fossils discovered in Africa, Europe, Asia, and South America (Lee 1997).

Pareiasaurs were common primary consumers in several terrestrial tetrapod faunas (Bernardi *et al*. 2017), including the late Permian fauna of North China (Sun 1980). Since 1960s, eight Chinese pareiasaur species have been described, including *Shihtienfenia permica* and *Shihtienfenia completus* from Baode, Shanxi (Young and Yeh 1963; Wang *et al*. 2019); *Shansisaurus xuecunensis*, *Huanghesaurus liulinensis*, and *Sanchuansaurus pygmaeus* from Liulin, Shanxi (Cheng 1980; Gao 1983, 1989); *Honania complicidentata* and *Tsiyuania simplicidentata* from Jiyuan, Henan (Young 1979); and *Elginia wuyongae* from Baotou, Inner Mongolia (Liu and Bever 2018). Among these species, only *Shihtienfenia completus* is based on a complete skull, the other species were established based on either postcranial materials or incomplete skulls or mandibles. Therefore, it is not surprising that the taxonomy and phylogenetic relationships of Chinese pareiasaurs are controversial.

Because some Chinese species were established based on different regions of the body and cannot be directly compared with each other, it is challenging to determine how many species are valid. Previously, different researchers proposed differing opinions; Lee (1997) argued that the teeth of *Honania* and *Tsiyuania* belong to the upper and lower jaws of the same individual and declared both to be *nomen invalid*. Lee also referred *Huanghesaurus* to *Shansisaurus*, while Li and Liu (2013) suggested that *Sanchuansaurus* and *Huanghesaurus* should be junior synonyms of *Shansisaurus*. After studying additional specimens from the type locality, Xu *et al* (2015) confirmed that the teeth previously attributed to *Honania* and *Tsiyuania* belong to the upper and lower jaws of the same species and resurrected the name *H. complicidentata*. Benton (2016) retained the dwarf-sized *Sanchuansaurus* and referred *Shansisaurus* and *Huanghesaurus* to *Shihtienfenia*.

In 2015, numerous disarticulated specimens were excavated from the Sunjiagou Formation in Baode, Shanxi (Dong and Yi 2018). Among these specimens, the first complete skull of Chinese pareiasaurs was described and tentatively named as *Shihtienfenia completus* (Wang *et al*. 2019). In 2017, the fossil hunter discovered some bones of pareiasaur and dicynodont specimens from both the Sunjiagou Formation and Upper Shihhotse Formation in Yangquan, Shanxi (Yi and Liu 2020). In 2018, a new pareiasaur skeleton (IVPP V33181) was collected from the Naobaogou Formation in Nei Mongol, China (Fig. 1A). The new specimen exhibits several features based on which a new species can be established. A phylogenetic analysis supports that IVPP V33181 and *S. completus* form a monophyletic clade, which is considered the sister group of Pumiliopareiasauria.

**FIG. 1.**
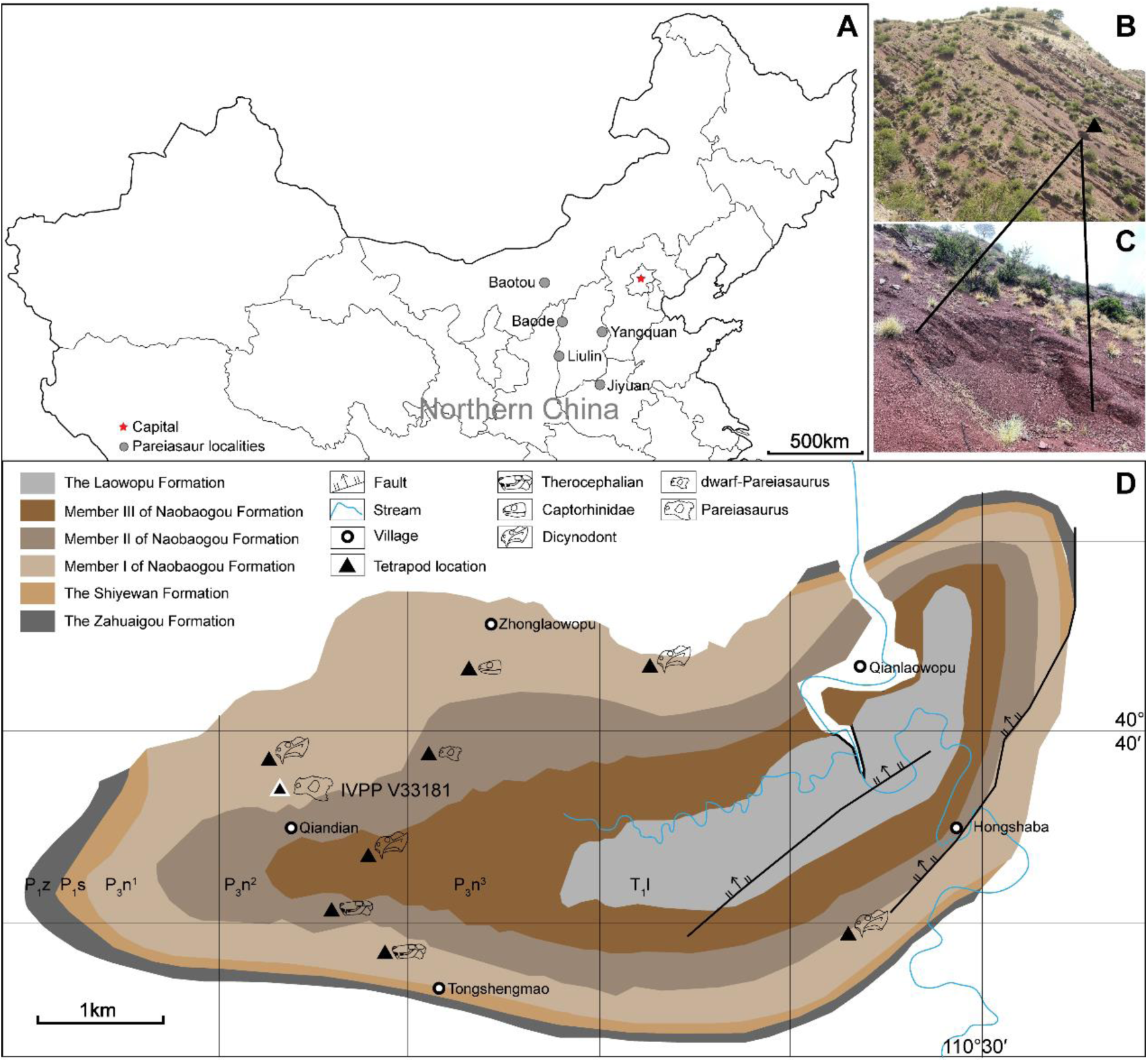
A, geographical location of Daqing Mountains fossil localities (dots). B, C, fossil locality photos of IVPP V 33181 (black triangle). D, geological map showing the tretrapod fossils (black trangles). *Abbreviations*: P_1_z, Zahuaigou Formation; P_1_s, Shiyewan Formation; P_3_n^1^, Member I of Naobaogou Formation; P_3_n^2^, Member II of Naobaogou Formation; P_3_n^3^, Member III of Naobaogou Formation; T_1_l, Laowopu Formation.

## GEOLOGICAL SETTING

IVPP V 33181 was excavated from purplish silty mudstone in the upper part of Member I of the Naobaogou Formation (Fig. 1B, 1C), near Qiandian Village, Shiguai District of Baotou City, Nei Mongol, China (Fig. 1D). Previously, Member I of the Naobaogou Formation yielded the dicynodont *Daqingshanodon limbus* (Zhu 1989) and *Jimusaria monanensis* (Shi and Liu 2023), the therocephalian *Euchambersia liuyudongi* (Liu and Abdala 2022), and the captorhinid *Gansurhinus naobaogouensis* (Liu 2023).

*Institutional abbreviations*. CAGS, Chinese Academy of Geological Sciences, Beijing, China; IVPP, Institute of Vertebrate Paleontology and Paleoanthropology, Chinese Academy of Sciences, Beijing, China; NMNH, Inner Mongolia Museum of Nature History, Hohhot, China; PIN, Paleontological Institute, Moscow, Russia; SXNHM, Shanxi Natural History Museum, Taiyuan, China.

## MATERIAL AND METHOD

### Phylogenetic analysis

We test the phylogenetic position of IVPP V 33181 using the data matrix modified from that of Van den Brandt *et al*. (2023), which comprises 142 characters. Two characters are revised in this study.

Character 54 (Postorbital region of skull, length) is revised as: **length at least equals preorbital region of the skull (0); postorbital region shorter than preorbital region (1).**

Character 56 (frontal, shape) is revised as: **slim and long, four times as long as wide (0); frontal length reduced, three times as long as wide (1); frontals short, with a length not more than two times the width (2).**

The coding for most taxa in Van den Brandt *et al*. (2023) are adopted, while *S. completus* and IVPP V33181 are included in this analysis. All 142 characters are analyzed as unordered and equally weighted. The heuristic maximum parsimony analysis, conducted using TNT 1.5, included 1000 addition sequences (repls.) and TBR branch swapping with 10 trees retained per replication. The default setting of collapsing zero-length branches was maintained. The search found 210 most parsimonious trees (MPTs) of length 298 steps with a retention index (RI) of 0.752 and a consistency index (CI) of 0.564).

### Nomenclatural acts

The nomenclatural acts has been registered in ZooBank, the online registration system for the International Commission on Zoological Nomenclature (ICZN). The ZooBank LSIDs (Life Science Identifiers) for this publication is: urn: lsid: zoobank.org: act: ADFF82E1-FBFE-448C-8268-F7BC5FD56880.

## SYSTEMATIC PALAEONTOLOGY

PAREIASAURIA Seeley, 1888

*Yinshanosaurus angustus* gen. et sp. nov.

*Holotype*. IVPP V 33181, an articulated skeleton with a nearly complete skull, paired scapulocoracoids, articulated 3rd-5th cervical vertebrae and 16 dorsal vertebrae, an incomplete ilium.

*Etymology*. ‘Yinshan’, one ancient name of the mountain where the specimen was collected; ‘*angustus*’, ‘narrow’ in Latin.

*Diagnosis*. *Yinshanosaurus angustus* and *Shihtienfenia completus* can be differentiated from all other pareiasaurs by the following combination of characters: (1) prominent postfrontal horn present; (2) pineal foramen situated about halfway along interparietal suture; (3) paired tubulars meet in midline, excluding postparietal from posterior margin of skull roof (also in Elginiidae); (4) skull with a postorbital region shorter than preorbital region.

*Yinshanosaurus angustus* can be distinguished from *Shihtienfenia completus* by the following features: (1) skull high and narrow, skull table longer than wide between two quadratojugals; (2) snout dimension as high as wide (3) length/width ratio of the frontal large (∼ 3.0); (4) U-shaped paraoccipital process in occipital view; (5) vertically oriented maxillary teeth.

### Description

*Skull table*. The skull is eroded and crushed, so the left cheek is heavily damaged while the right cheek is nearly intact but slightly laterally everted. Additionally, the skull roof was somewhat twisted (Fig. 2). The skull measures approximately 37 cm from the anterior tip of the premaxilla to the posterior margin of the tabular along the dorsal midline, which is comparable to the skull length of *Shihtienfenia completus* (IMMNH-PV00020) (Wang *et al*. 2019). The length of skull table is greater than the width between the quadratojugals. The outer surface bears blunted bosses and sparse roughly circular pits. Ridges radiated on skull as in *Deltavjatia* (Tsuji 2013) and *Pareiasuchus* (Lee and Kitching 1997), similar ridges are absent in *Anthodon* (Lee 1997). Pairs of small central bosses are located on the frontal and parietal, respectively, which are common in many pareiasaurs other than *Elginia*, *Arganaceras*, *Pareiasaurus* and *Bunostegos* (Tsuji *et al*. 2013). Two pairs of bosses are prominent on the nasal and the postfrontal, but such bosses are absent on the supratemporal. The nasal bosses are rounded, and the postfrontal bosses are blunted, differing from the pointed postfrontal horns in *S. completus* (Wang *et al*. 2019).

**FIG. 2.**
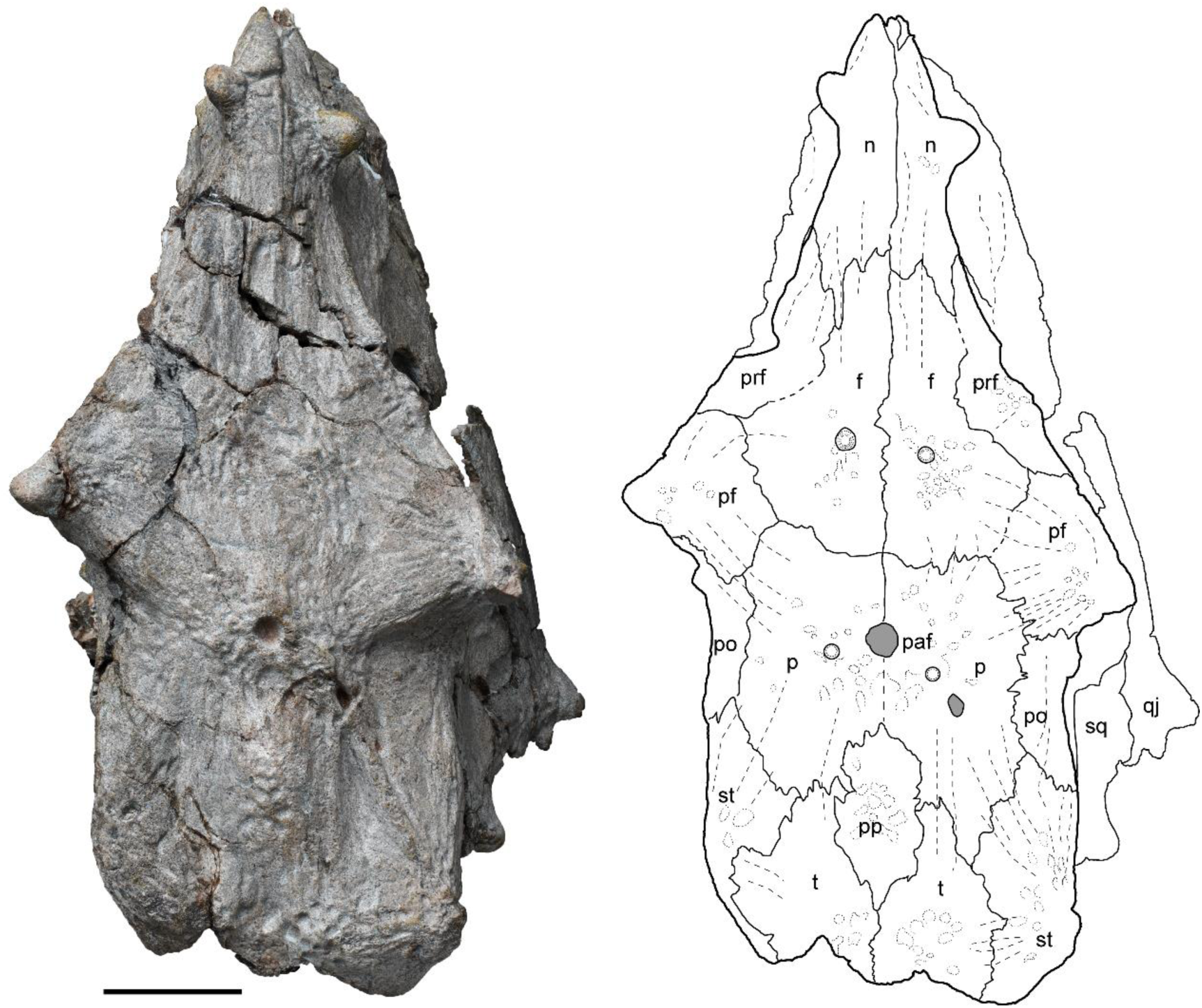
Skull of *Yinshanosaurus angustus* sp. nov., holotype IVPP V33181 in dorsal view. *Abbreviations*: f, frontal; n, nasal; p, parietal; paf, prietal foramen; pf, postfrontal; po, postorbital; pp, postfrontal; prf, prefrontal; qj, quadratojugal; sq, squmosal; st, supratemporal; t, tabular. Scale bar represent 50mm.

The long **frontal** is a roughly pentagonal bone on the skull roof (Fig. 2). It meets its counterpart along a nearly straight suture on the midline. It sutures with the nasal anteriorly, the prefrontal anterolaterally, the postfrontal posterolaterally, and the parietal posteriorly. The frontal is excluded from the orbital margin due to the contact between the prefrontal and the postfrontal. On the anterior portion, the dorsal surface bears several longitudinal ridges, including the midline one, and depressions. The dorsal surface displays a prominent central boss, on the rest, there are a cluster of smaller bosses close to the midline, at the junction where the frontal, prefrontal and postfrontal meet. The frontals contribute to the anterior portion of the skull table, which is controlled by the width of the skull. The length/width ratio of frontal in *Y. angustus* (3.0) is significantly larger than that of the other pareiasaurs (2.0).

The **prefrontal** constitutes the anterodorsal portion of the orbital rim (Fig. 3). It features three interconnected bosses (one large and two small) along the lateral edge. On the skull roof, the prefrontal is a narrow bone lateral to the frontal. It contacts the nasal anteromedially and the postfrontal posteriorly.

**FIG. 3.**
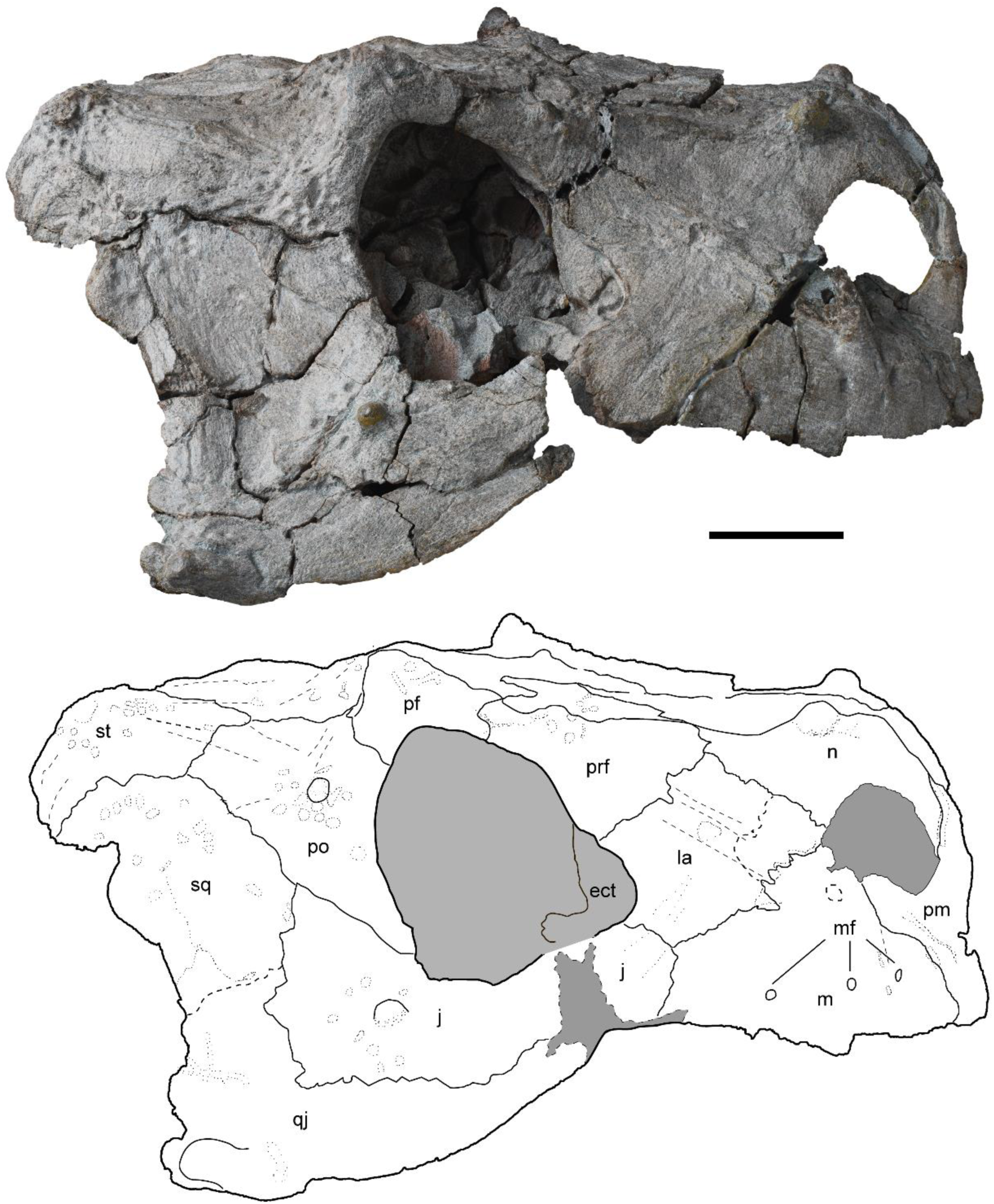
Skull of *Yinshanosaurus angustus* sp. nov., holotype IVPP V33181 in lateral view. *Abbreviations*: ect; ectopterygoid; f, frontal; j, jugal; l, lacrimal; m, maxilar; mf, maxilar foramen; n, nasal; pf, postfrontal; pm, premaxilar; po, postorbital; pp, postfrontal; prf, prefrontal; qj, quadratojugal; sq, squmosal; st, supratemporal. Scale bar represent 50mm.

In lateral view, the **postfrontal** is a curved bone, defining the dorsal rim of the orbit.

It has a short suture with the prefrontal anteriorly, a long suture with the postorbital posteroventrally. On the skull roof, it meets the frontal anteromedially and the parietal posteromedially. A large, blunt-pointed horn is situated at the midpoint of the orbital rim, pointing dorsolaterally, while several ridges radiate outwards from this location medially.

Pareiasaurs’ skulls are ornamented by multiple bosses, the prominent pairs of postfrontal horns are well developed in *Elginia* (Newton 1893; Liu and Bever 2018), *Arganaceras* (Jalil and Janvier 2005) and *S*. *completus* (Wang *et al*. 2019).

Only the right **premaxilla** is preserved on the anterior edge of the skull (Figs. 3, 4). It is a triradiate element, exposed in lateral, medial, and ventral views. It has a long, slender dorsal process which inserts between the two anterior processes of the nasals and forms the medial/anterior rims of the naris, and a lateral process that sutures with the anterior margin of the maxilla. Laterally, the premaxilla-maxilla suture lies above the third tooth alveolus, indicating that it should accommodate three teeth in total. On the palate, the bone sutures with the maxilla posterolaterally and contacts the anterior most portion of the vomer posteriorly. The prepalatal foramen cannot be observed due to the absence of the left premaxilla.

The **maxilla** features a relatively smooth lateral surface (Fig. 3). It extends anteriorly and slightly overlaps the lateral surface of the premaxilla. It sutures with the lacrimal posterodorsally, the jugal posteriorly. Its anterodorsal process is swollen but is broken for the upper portion, it is unknown whether there was a boss or a horn as in *Elginia*, *Scutosaurus* or *Sanchuansaurus*. The anterodorsal process contributes to the lateral/posterior rim of the naris. On the lateral surface, the posterior end of the maxilla lies anterior to the orbit, but it still extends posteriorly medial to jugal behind the anterior rim of the orbit. It is unknown whether the maxilla contacts the quadratojugal, while this contact is typically regarded as a synapomorphy of pareiasaurs (Lee 1997). The maxilla bears three distinct foramina dorsal to the ventral (teeth-bearing) margin, which probably transmitted fibers of the maxillary branch of the trigeminal nerve or as the artery canal (Tsuji *et al*. 2013). The distance between the two closely spaced anterior foramina is half the length of the distance between the two posterior ones. On the palatal surface, the maxilla forms the lateral border of the choana, and sutures with the palatine and the ectopterygoid posteromedially (Fig. 4). The maxilla bears at least 13 marginal tooth alveoli, however, there are only 9 functional teeth remaining.

**FIG. 4.**
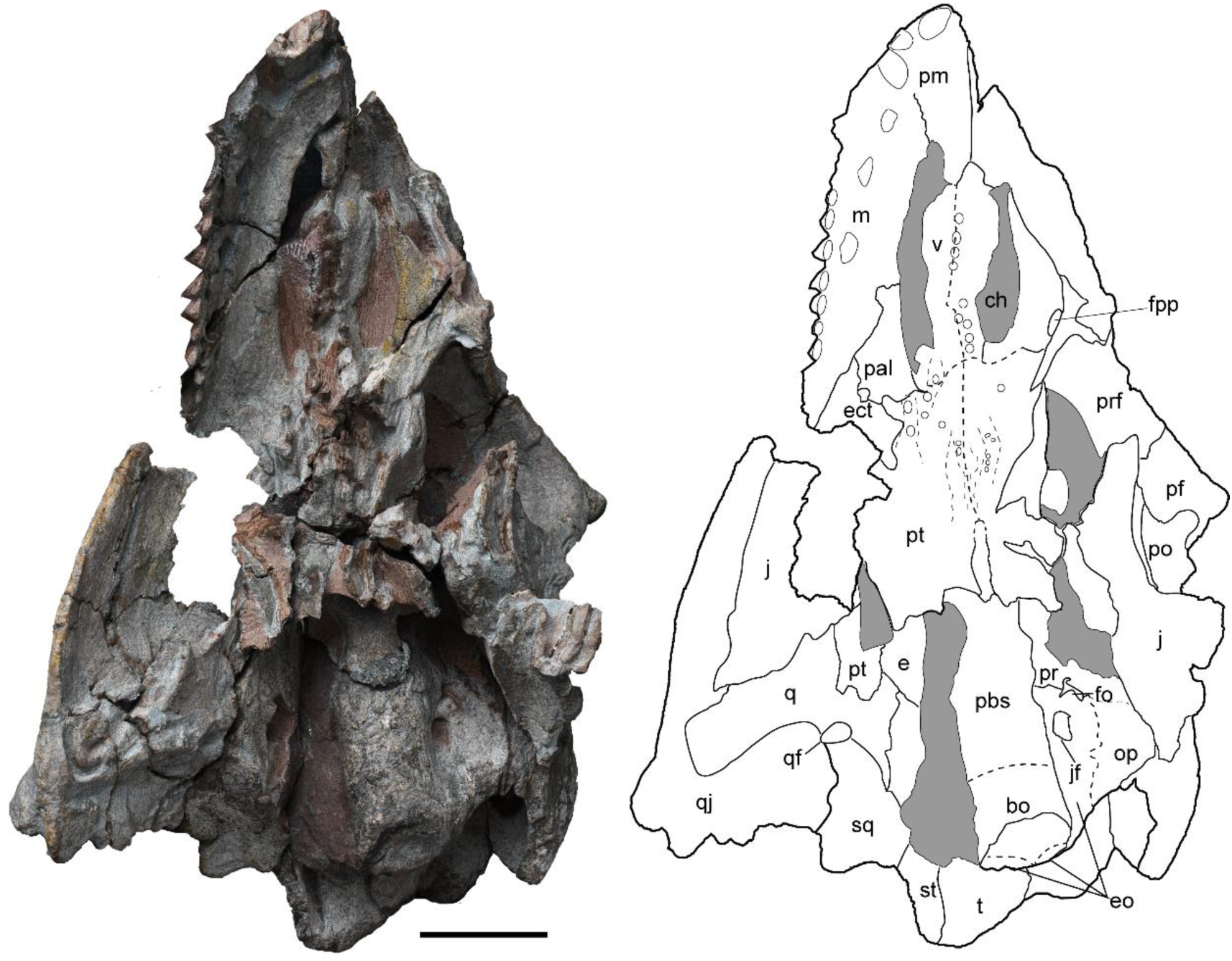
Skull of *Yinshanosaurus angustus* sp. nov., holotype IVPP V33181 in ventral view. *Abbreviations*: bo, bosioccipital; ch, choana; e, epipterygoid; ect; ectopterygoid; eo, exoccipital; f, frontal; fo, fenestra ovalis; fpp, foramen palatinum posterius; j, jugal; jf; foramen jugulare anterius; m, maxilar; op, opisthotic; pal, palatal; pbs, parabasiosphenoid; pf, postfrontal; pm, premaxilar; po, postorbital; pp, postfrontal; pr, prootic; prf, prefrontal; pt, ptergoid; q, quadrate; qj, quadratojugal; sq, squmosal; st, supratemporal; t, tabular; v, vomer. Scale bar represent 50mm.

The **lacrimal** is roughly quadrilateral between the naris and the orbit (Fig. 3). It contributes to the posterior corner of the naris and the anterior border of the orbit. It sutures with the nasal anterodorsally, the prefrontal posterodorsally, and the jugal posteroventrally. It has a slanted, wavy suture with the maxilla anteroventrally. Its posterior portion contacts the ectopterygoid medially to the orbital margin. A ridge, which begins from the maxillary dorsal process and ends on the anterior border of prefrontal, runs posterodorsally along the middle lateral surface of the lacrimal, dividing the bone into dorsal and ventral portions.

The **nasal** is a roughly triangular bone in dorsal view and pinches in the height in lateral view (Figs. 2, 3). It has a tapering anterior process to receive the premaxillary dorsal process. The nasal meets its counterpart along the midline, and the frontal posteriorly with serrated suture. On the lateral surface, it sutures with the lacrimal posteroventrally and the prefrontal posteriorly. This bone forms the upper half rim of the naris. The rounded nasal boss, located above the posterior rim of the naris, is oriented dorsolaterally. The large, rounded naris opens mainly anteriorly, resembling the structure found in *S. completus* (Wang *et al*. 2019). The premaximal, lacrimal, maximal, and nasal form the border of snout. Anterodorsally, the snout dimensions in *Bunostegos*, *Pareiasuchus* (Tsuji *et al*. 2013), and *S. completus* (Wang *et al*. 2019) is broader, whereas in *Arganaceras*, *Elginia* (Jalil and Janvier 2005) and *Yinshanosaurus*, the narrow skull makes the snout as high as wide.

*Cheek*. The lateral surface of the cheek consists of the postorbital, the jugal, the squamosal, and the quadratojugal (Fig. 3). The posterior margin of the cheek is opisthocoelous in *S. completus* and *Y. angustus*, so the postorbital region consisting of the squamosal and quadratojugal is shorter than the preorbital region consisted by the premaxilla, the nasal, the maxilla and the lacrimal. This feature is unique in *Arganaceras* (Jalil and Janvier 2005), *S*. *completus* (Wang *et al*. 2019) and *Y. angustus*, that differs from other pareiasaurs.

The **postorbital** is subtriangular in shape (Fig. 3). Its anterior edge forms the posterior rim of the orbit. It sutures with the postfrontal anterodorsally, the parietal dorsally, and the supratemporal posterodorsally. Its posteroventral margin sutures to the squamosal and jugal. A prominent hump surrounded by a cluster of pits is located at the middle of the bone.

The **jugal** is boomerang-shaped, forming the ventral rim of the orbit (Fig. 3). Its anterior process laterally overlaps the posterior process of the maxilla and the posteroventral corner of the lacrimal. Its ventral portion covers the upper part of the quadratojugal, while the posterior margin sutures with quadratojugal and squamosal. It has a relatively smooth lateral surface, except for a single boss surrounded by several pits, which is located slightly posterior to the orbit. This boss is slightly larger than the one on the postorbital.

The **squamosal** forms the upper part of the cheek (Fig. 3). This element appears pentagonal in lateral view. It sutures with the jugal anteroventrally, the postorbital anterodorsally, the supratemporal dorsally, and the quadratojugal ventrally. It has an irregular posterior margin. Its lateral surface is sculptured by patchy pits. On the posterior edge, a large oblong boss points backwards. Interiorly, the squamosal articulates with the quadrate, forming a vertical wall that supports the cheek. The upper portion of the squamosal contributes to the articulation that connects to the paraoccipital process.

The L-shaped **quadratojugal** forms the entire ventral and lower posterior margin of the cheek (Fig. 3). It delineates most of the ventral and posterior boundaries of the jugal, and dorsally, it forms a relatively straight horizontal suture with the squamosal. Two pointed bosses are located at the posterior corner of the quadratojugal. The anterior boss directs posterolaterally, while the posterior one directs posteriorly. Above these, an oblong boss extends backwards along the posterior edge. In occipital view, the quadratojugal sutures with the quadrate ventrally (Fig. 5), rendering the quadrate joint invisible in lateral view.

**FIG. 5.**
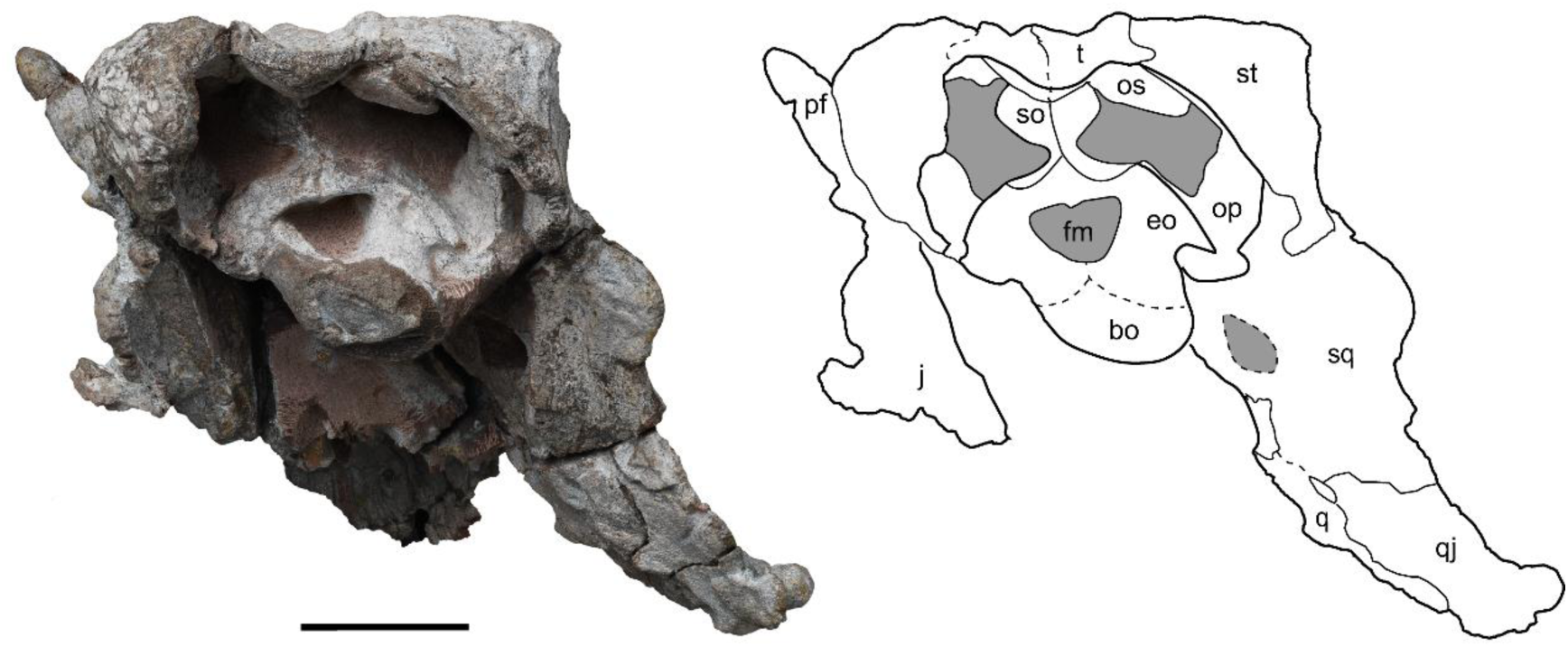
Skull of *Yinshanosaurus angustus* sp. nov., holotype IVPP V33181 in occipital view. *Abbreviations*: bo, bosioccipital; eo, exoccipital; fm, foramen magnum; j, jugal; op, opisthotic; os, osteoderm; q, quadrate; qj, quadratojugal; sq, squmosal; so, supraoccipital; st, supratemporal; t, tabular. Scale bar represent 50mm.

The **parietal** is a large, pentagonal bone, which is isometrical in both length and width relative to the frontal (Fig. 2). It has a slightly zigzag lateral suture with the postfrontal and postorbital. Its posterior margins form an acute posterior process, inserting between the supratemporal and the tabular. The rounded pineal foramen is located on the midpoint of the suture between the paired bones. The area surrounding the pineal foramen is elevated. The pineal foramen is the external opening of pineal gland which is located between the cerebral hemispheres. In the midline of the skull roof, the parietal and postparietal lie anteroposteriorly, the parietal became shorter when the postparietal move forward, the position of pineal foramen is corresponding backward.

On the right parietal, there is an almond-like hole surrounded by swelled margin, possibly a healing bite mark inflicted by a carnivore (therocephalian).

The single **postparietal** is a hexagon bone, raised as a median bulge on the midline of the skull roof (Fig. 2). It sutures with the paired parietals anteriorly and is surrounded by the paired tabular posteriorly, thereby excluding it from the posterior margin of the skull roof. The dorsal surface of the postparietal is heavily sculptured by a cluster of pits. In occipital view, the postparietal extends ventrally to form a short vertical mid-ridge, which connects to the dorsal process of the supraoccipital and forms the primary dorsal hinge between the braincase and the skull roof (Fig. 5).

The **tabular** is a small pentagonal bone (Fig. 2). The suture between the paired bones constitutes the terminal midline of the skull roof. Their posterior margins are interrupted by notches. The dorsal surface is sculptured by a cluster of pits located on the posterior part. In ventral view, the tabular extends laterally to reinforce the skull roof, which makes an imbricate contact between the tabular and the supratemporal (Fig. 4).

Tabular was the ‘supernumerary element’ interposed between the postparietal and supratemporal (Lee 1997), it was homologous to the tabular in parareptiles such as *Macroleter* and *Nyctiphruretus*(Tsuji 2010). The paired tubulars were separated by the interposed postparietal in many pareiasaurs, only contact in the specialized Elginiidae. The curved **supratemporal** forms the posterolateral corner of the skull roof (Figs. 2, 3). On the skull roof, it contacts the postorbital anteriorly, the parietal anteromedially, and the tabular medially. Ventrally, it sutures with the squamosal (Fig. 3). The left boss is worn, while the right one is fractured, only the base is preserved. Three series of ridges radiate outward from the base of the boss, while the remaining part is sculptured by a cluster of pits.

*Palate*. The palate is characterized by several denticle-bearing ridges and two kidney-shaped choanae. Many of the bone sutures on the palate are unclear and are indicated by dotted lines (Fig. 4).

The **vomer** is a long, elliptical bone (Fig. 4). It sutures with the premaxilla anteriorly, the palate posteriorly, and meets its counterpart at the midline. Nine denticles arranged in a single row lie along the median edge of the left vomer, with the caniniform cusp of each denticle curved posteroventrally.

The **palatine** is an irregularly polygon-shaped bone, forming the posterolateral edge of the choana (Fig. 4). It sutures with the maxilla anterolaterally and the ectopterygoid posterolaterally. Its medial margin sutures with the pterygoid. There is no denticle on this bone. The foramen palatinum posterius (suborbital) is located between the palatine and the ectopterygoid.

The small **ectopterygoid** sutures with the palatine anteromedially, the pterygoid posteromedially, and the maxilla laterally (Fig. 4). Its posterior margin contributes to the anterior margin of the subtemporal fossa. The ventral surface is smooth. In lateral view, the dorsal process of the ectopterygoid constitutes the anterior inner rim of the orbit.

The **pterygoid** forms the posterior portion of the palate, featured by a series of denticles (Fig. 4). It sutures with the vomer anteriorly, the palatine and ectopterygoid anterolaterally, and the quadrate posterolaterally. It meets below the parabasisphenoid rostrum and its counterpart at the midline. The interpterygoid vacuity is a narrow slit with a broken anterior margin. The transverse flange is posteroventrally deformed and lacks denticles. The narrow flange indicates weak contact between the pterygoid and the cheek. The pterygoid has a wide quadrate process. Its distal portion is flared and forms a serrated suture with the quadrate, while the dorsal portion extends over the dorsal flange of the quadrate and contacts the squamosal posterodorsally. Multiple rows of denticles traverse the ventral surface of the pterygoid. The paired medial rows of denticles extend from the vomer and close to the margin of the interpterygoid vacuity. Other rows of denticles occupy the palatine branch.

The left **quadrate** is broken, while the right is complete (Fig. 4). Dorsally, the quadrate has an expanding flange. It forms an extensive suture with the quadrate flange of pterygoid anteromedially, sutures with the quadratojugal at its ventral and posteroventral margins, and the squamosal at its posterodorsal margin. The quadrate foramen lies at the convergence among the quadrate, squamosal, and quadratojugal. It forms an interlayer with the cheek anteriorly and is fused with the quadratojugal and squamosal posteriorly. This unique structure provides an adequate surface to attach the masseter muscle. The quadrate connects the palate with the cheek. The quadrate body is dorsoventrally expanded and is nearly perpendicular to the basisphenoids. Ventrally, the quadrate consists of a lateral and a medial condyle which are anteroposteriorly enfolded.

*Braincase*. In most pareiasaur specimens, the elements of the braincase are fused. In the new form, the braincase is also well-ossified, making the sutures difficult to identify in the ventral view. Nevertheless, additional morphological details can be observed through the exposure of the right lateral portion of the braincase (Fig. 3).

The basisphenoid and parasphenoid are fused to form the **parabasisphenoid** (Fig. 4). The anterior end of this bone is bifurcated into two basipterygoid processes, which are not fused to the pterygoid. It is constricted in the neck and has a cylindrical shape in the posterior portion, lacking a distinct basal tubercle. The smooth ventral surface features two lateral crests that extend from the middle of the bone to the anterior part of the basioccipital.

In occipital view, the **occipital condyle** is trichotomous, formed by the basioccipital ventrally and the exoccipitals laterally (Fig. 5). The basioccipital portion is incomplete due to erosion. Additionally, there is no distinct suture between the basioccipital and the parabasisphenoid or the exoccipitals.

The paired **exoccipital**s converge at the midline, forming the ventral and lateral margins of the foramen magnum (Fig. 5). The dorsal portion of the exoccipital is crescent and oblique dorsolaterally, while its medial part provides support for the supraoccipital. The lateral process exhibits a long suture with the paraoccipital process. On the ventrolateral side, the dorsolateral projection of the exoccipital delineates two large foramina: the fenestra ovalis and the foramen jugulare anterius (Fig. 4). The anteriorly located fenestra ovalis is smaller than the posteriorly situated foramen jugulare anterius. The foramina for cranial nerves IX-XII are indistinct.

The **supraoccipital** connects the skull roof to the braincase (Fig. 5). In occipital view, this element is hourglass-shaped, narrowing at the waist to form a posteriorly oriented sagittal crest. The supraoccipital also constitutes the medial wall of the post-temporal fenestra.

The left **opisthotic** is almost exposed in ventral view (Fig. 4). It articulates with the prootic anteriorly, the supratemporal laterally, and the exoccipital posteriorly and medially. On the occiput, the paraoccipital process extends laterally and expands dorsoventrally, forming a ‘U’-shaped dorsal margin that forms the ventral border of the post-temporal fenestra. The lateral margin articulates broadly with the squamosal, while its curved dorsal corner makes brief contact with the supratemporal. The paraoccipital process is the connection between the posterior portion of the cheek and the braincase and reaches to the posterior margin of the skull roof (Fig. 4). The lateral eversion of the cheek determines the orientation of the paraoccipital process. The process is restrained forming a U-shape as in the African pareiasaurs such as *Embrithosaurus*, *Nochelesaurus*, and *Bradysaurus* (Van den Brandt *et al*. 2023), *Bunostegos* (Tsuji *et al*. 2013).

The left **prootic** is exposed on the anterolateral side of the braincase (Fig. 4). It is fused with the parabasisphenoid ventrally and forms the anterior portion of the dorsal border of the foramen ovalis. Posteriorly, the prootic merges with the occipital portion of the opisthotic (the paraoccipital process).

The anterior portion of the braincase, including the right prootic, sphenethmoid, and epipterygoid is well exposed in the right orbit (Fig. 6).

**FIG. 6.**
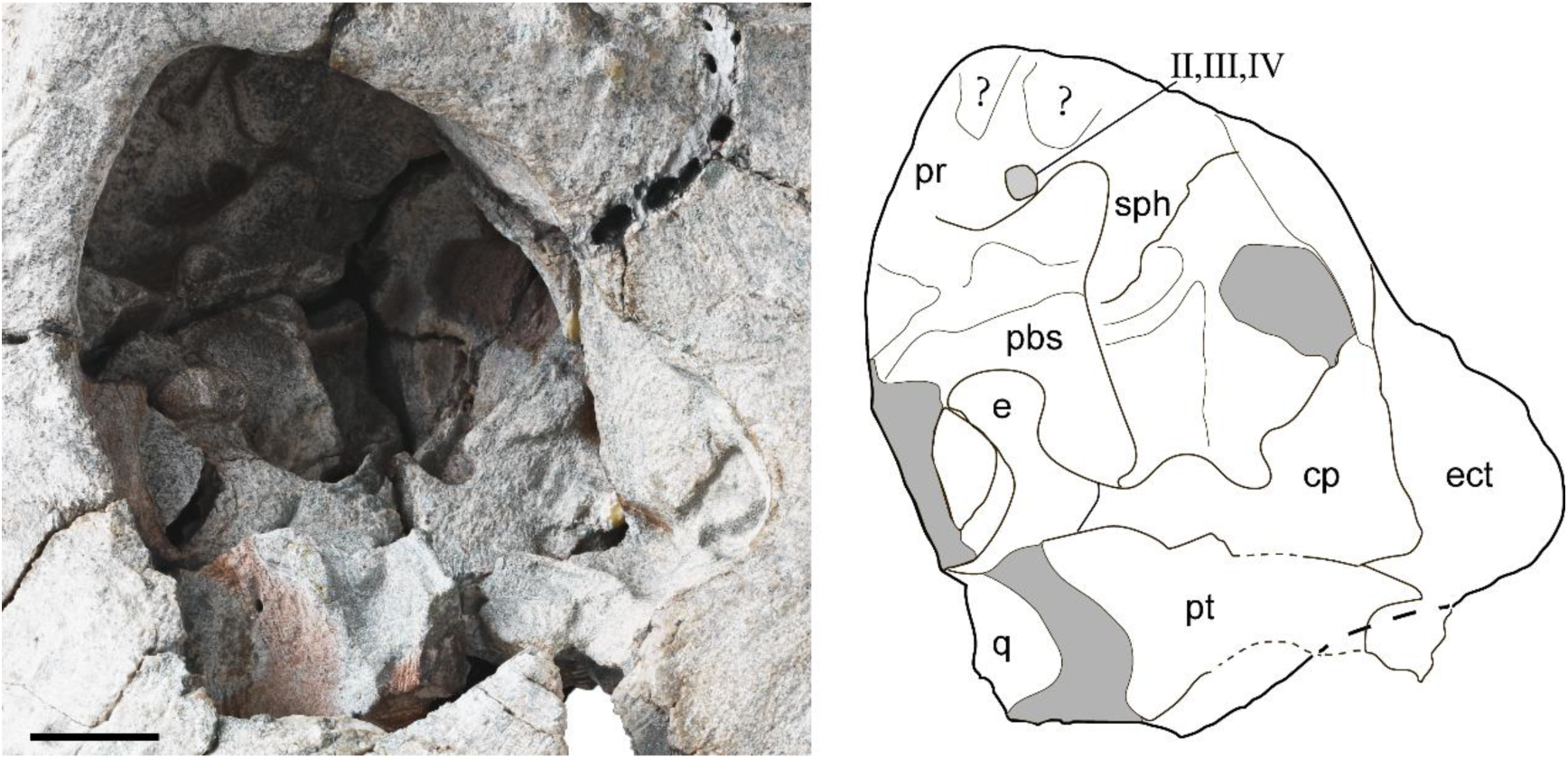
Anterior braincase of *Yinshanosaurus angustus* sp. nov., holotype IVPP V33181 in lateral view. *Abbreviations*: cp, cultriform process; e, epipterygoid; ect; ectopterygoid; pbs, parabasiosphenoid; pr, prootic; pt, ptergoid; q, quadrate; sph, sphenethmoid; II,optic nverve; III, oculomotor nverve; IV, trochlear nverve. Scale bar represent 20mm.

The well-ossified **sphenethmoid** is supported by the prootic and the cultriform process of the parabasisphenoid and locates beneath the frontal (Fig. 6). The optic, oculomotor, and trochlear nerves traverse together an adequate-sized foramen situated between the sphenethmoid and the prootic. The sphenethmoid has been described in *Bunostegos* and *Deltavjatia* (Tsuji 2013; Tsuji *et al*. 2013).

The **epipterygoid** sutures with the quadrate posteriorly, the pterygoid lateroventrally, and the cultriform process anteroventrally (Fig. 6). It covers the parabasisphenoid laterally and adheres to the postorbital medially. The clavate dorsal end of the epipterygoid process swells, which is supposed to be the junction with the oculomotor muscle or cartilage (Shi and Liu 2023).

*Dentition.* Only nine marginal teeth and one replacement tooth are exposed on the maxilla in lingual view (Fig. 7). The teeth are close-packed, and the triangle-shaped crown is slightly curved lingually. The roots of the teeth are noticeably constricted. On the replacement tooth, at least 7 cusps are evenly arranged along the margin of the tooth crown. The number of cusps was considered as an important distinguishing feature by Lee (1997). However, this factor probably indicates different ontogenetic stages of individuals.

**FIG. 7.**
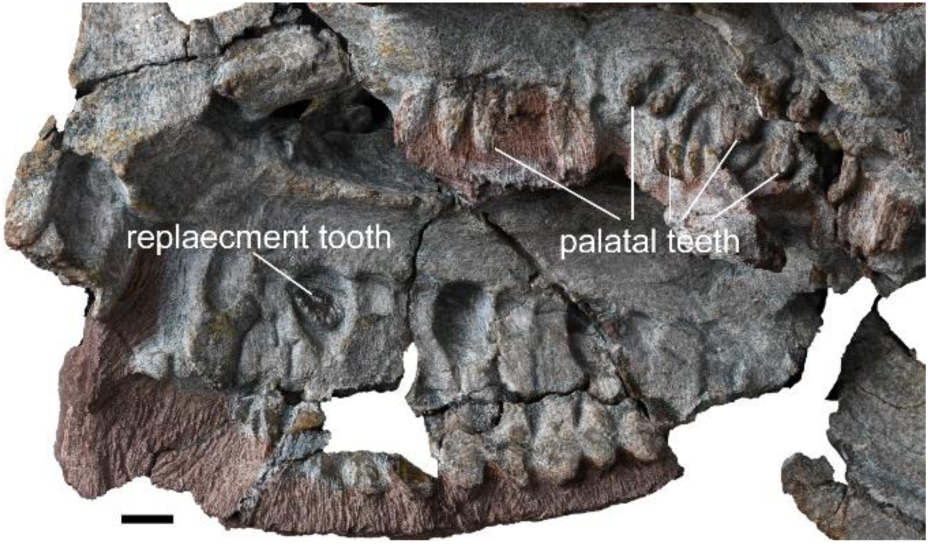
Skull of *Yinshanosaurus angustus* sp. nov., holotype IVPP V33181 in medial and left lateral view. Scale bar represent 10mm.

The occlusion of the jaws indicates that the lower jaw is embedded in the upper jaw in *S*. *completus*. Its maxillary teeth bend ventromedially, and its alveolar ridge is separated by an obvious crease from the ventral margin of maxilla. On the labial side, the alveolar ridge of *Y. angustus* is flat with the maxilla, all the teeth are orientated directly downwards.

*Scapulocoracoid*. Both scapulocoracoids are preserved: the right one is nearly complete, while the left one is broken into three pieces and hard to be reconstructed (Fig. 8A, 8B, 8C, 8D). The scapular portion is typically a straight element; however, it is significantly deformed on the right one. The bone is curved, and the dorsal end is twisted 90° from the original position.

**FIG. 8.**
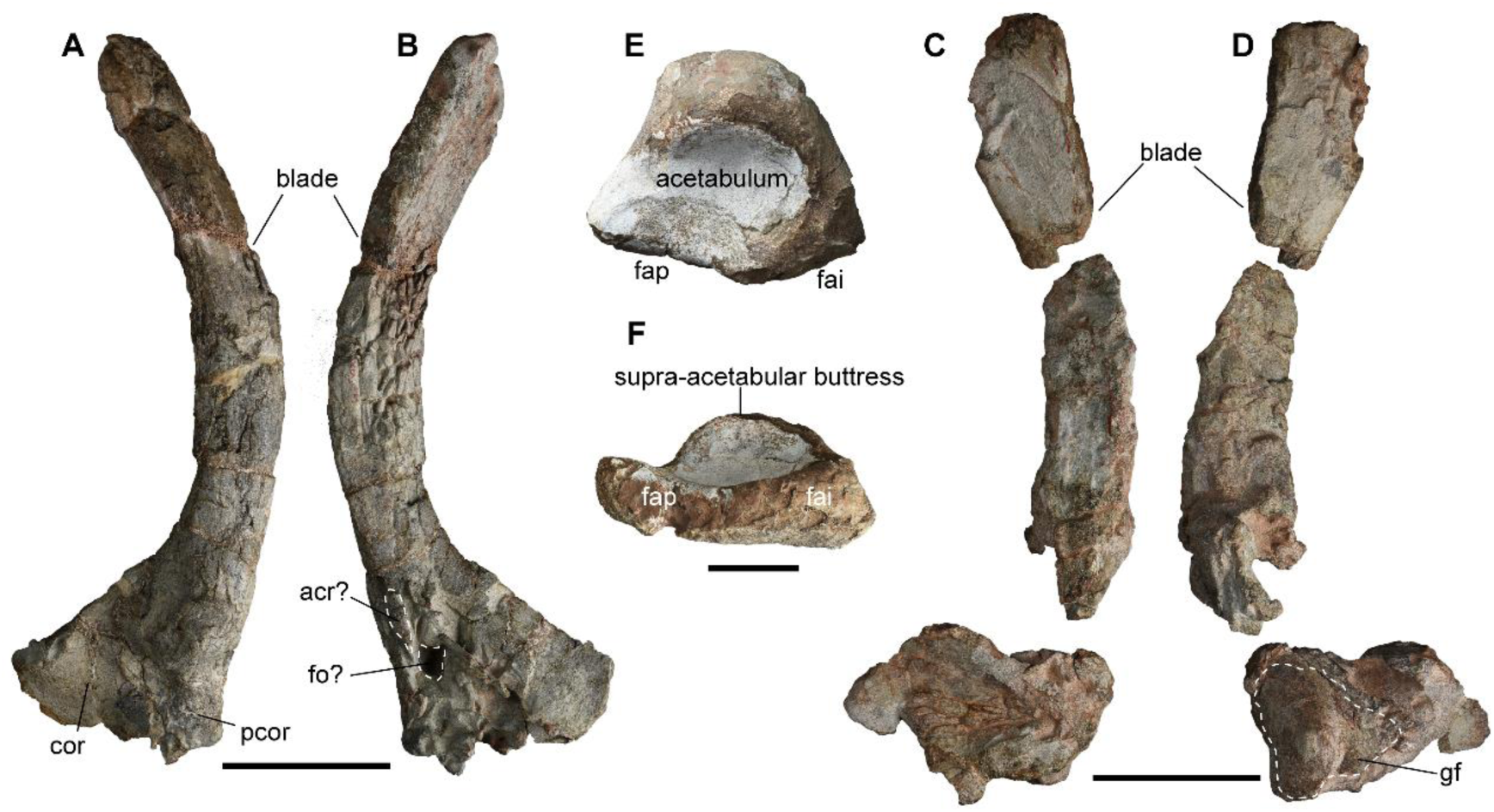
Girdle elements of *Yinshanosaurus angustus* sp. nov., Scapulocoracoid of holotype IVPP V33181 in lateral (A and D) and medial (B and C) view. Left ilium of holotype IVPP V33181 in left lateral (E) and ventral (F) view. Abbreviations: acr, acromoion process; cor, coracoid; fai, face articularis ischiatica; fap, face articularis pubis; fo, coracoid foramen; gf, glenoid fossa; pcor, procoracoid. Scale bar represent 100mm.

The left coracoid foramen is not observed on the lateral side, and the acromion process of the scapular portion is bent on the medial side, generating the glenoid fossa being unidentifiable. In lateral view, the scapular blade thins dorsoventrally. The distal portion of the blade is anisopleural, its anterior edge is angular, while the posterior edge is rounded. The suspected glenoid fossa is bipartite, with a short ridge separating the shallow fossa into two facets. The anterior facet is large and directed laterally, while the posterior facet is small and directed dorsoventrally.

The posterior and ventral margins of the coracoid portion form an angle of 45°. Several thickened coracoid rim buttresses are located along the margin of the coracoid. The procoracoid portion is fractured and bends toward the medial side. On medial view, the scapular portion is marked by striped nodules. The medial surface does not exhibit any identifiable characteristics.

Although the left scapulocoracoid is cracked, the glenoid can be identified. The curved shape of the scapular blade is confirmed due to the symmetrical element.

*Ilium* Only ventral portion of the left ilium is preserved (Fig. 8E, 8F). The supra-acetabular buttress of the ilium is well developed, forming a prominent tuberosity on the lateral side. Anterior to the acetabulum, there is a shallow groove with a rugose wall. In the center of the acetabulum, the acetabular fossa is clear. The dorsal outline of the acetabulum is approximately arc-shaped, with a notched front. The ventral side of the ilium consists of two facets for the pubis and ischium, forming a pointed angle anteriorly and a blunt margin posteriorly.

*Axial skeleton*. A series of 16 well-preserved vertebrae (3^rd^ to 18^th^) and three isolated incomplete vertebrae (19^th^ to 21^st^) are preserved. The first five of the series are presumed to belong to the cervical vertebrae based on the general characteristics of the vertebrae in pareiasaurs (Van den Brandt *et al*. 2021a). In the cervical region, both the parapophysis and the diapophysis form shallow facets on the centrum, whereas they are significantly expanded in the dorsal region. The cervical and dorsal columns are clearly distinguishable from one another.

Only the last two cervical vertebrae are well preserved, the third vertebra retains only the neural spine (Fig. 9A). A fragment of the osteoderm is preserved on the left lateral side.

**FIG. 9.**
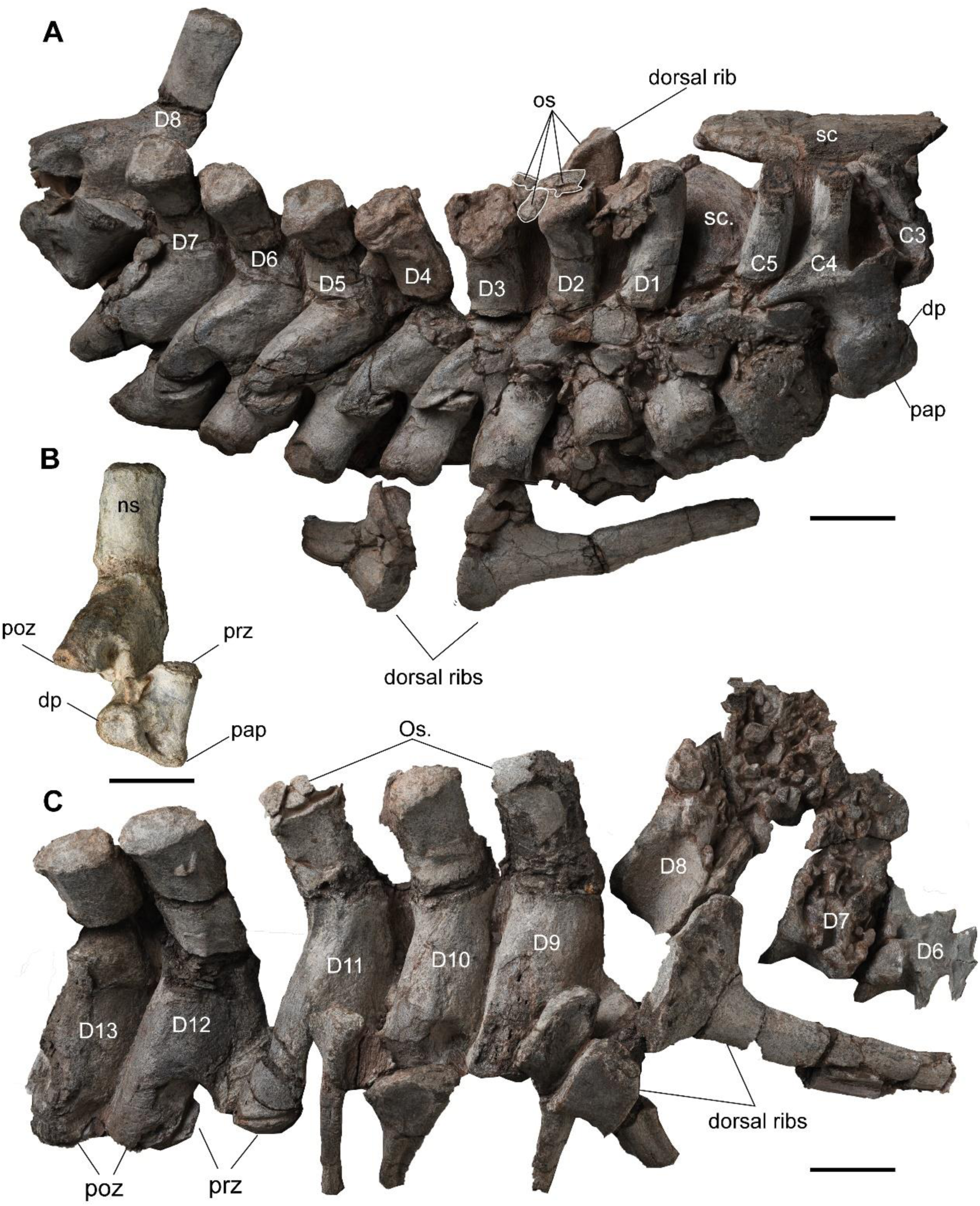
Cervical and dorsal columns of *Yinshanosaurus angustus* sp. nov. A, 3^rd^-5^th^ cervical and 1st-8^th^ dorsal vertebrae of holotype IVPP V33181 in right lateral view. B, 8^th^ dorsal vertebrae holotype IVPP V33181 in right lateral view. C, 6^th^-13^th^ dorsal vertebrae of holotype IVPP V33181 in right lateral view. Abbreviations: dp, diapophysis; ns, neural spine; os. osteoderm; pap, parapophysis; poz, postzygapophysis; prz, prezygapophysis; sc, scapula. Scale bar represent 50mm.

The fourth and fifth cervical vertebrae are well articulated. Compared to the dorsal vertebrae, the cervical vertebrae are taller than they are wide, featuring short and narrow pre- and postzygapophysis. The neural spine is tall and projects anterodorsally, with the distal portion expanding laterally to form a rectangular cross-section. The anteroposterior margin of the amphicoelous centrum is compressed, and the centrum appears rectangular in lateral view. Laterally, both the parapophysis and the diapophysis are worn down, leaving only a cross groove preserved between the two apophyses. The prezygapophysis is significantly smaller than the postzygapophysis. The small and short surface of the prezygapophysis is slightly bent medially, while the surface of the postzygapophysis curves lateroventrally. The postzygapophysis extends posterolaterally from the base of the neural spine, where a distinct crest extends to the postzygapophysis.

Thirteen articulated dorsal vertebrae (6^th^ to 18^th^) are exposed on the right side, while most centra are encased in the surrounding rock. Only the 8^th^ dorsal vertebra (13^th^) is well exposed (Fig. 9B); its centrum is narrow, taller than broad, with deeply concave sides. The face articularis develops considerably, featuring a large overlapping area between the pre- and postzygapophysis. In dorsal view, the distal ends of the neural spines gradually narrow lateromedially from the cervicals to the dorsals. In lateral view, the diapophysis projects downward from the cervicals to the dorsals in a gradual manner. The 6th to 13th vertebrae are preserved in a single block (Fig. 9A). The first two dorsal vertebrae (6th and 7th) retain some features of the cervical vertebrae (Fig. 9A), including an anterodorsally bending neural spine, narrow pre- and postzygapophysis, and pointed postzygapophysis. Nevertheless, the prominent parapophysis and diapophysis, which articulate with the dorsal ribs, distinguish these vertebrae from the cervical ones. Their neural spines are significantly more robust than those of the cervicals. Several contiguous osteoderms cover the dorsal facet of the neural spine. The outer surface of the osteoderms is rough, featuring tubercles, pits, ridges, channels, and openings. The ventral surface is convex, conforming to the concave surface of the neural spine. A middle ridge extends dorsoventrally along the anterior margin of the spine but does not reach the upper one-third portion. The parapophysis and diapophysis are separated by a groove, and the facet on the upper parapophysis is larger than that on the lower diapophysis. Both facets are oriented ventrolaterally.

The 8th to 13th vertebrae features similar. Fragments of osteoderm are present on the top of the neural spine. The neural spine is cylinder-shaped, and the anterior middle crest reaching its apex. The transverse processes expand laterally, forming a flange-like wall. Mediolaterally, the dorsal vertebrae are significantly wider than the cervicals, while anteroposteriorly, the dorsals are shorter than the cervicals. The pre- and postzygapophysis articulate tightly. The prezygapophysis is oriented dorsally, while the postzygapophysis is oriented ventrally; the groove between the parapophysis and the diapophysis is shallow. The diapophysis articulates with the parapophysis as one entirely surface in the eighth dorsal (13^th^) (Fig. 9B).

The 14^th^ to 18^th^ vertebrae are preserved in a separate block (Fig. 9C), with only the upper part of the right side exposed. The distal end of the neural spine narrows in the anteroposterior direction. Ridges are present on both the anterior and posterior margins of the spine.

Several disarticulated dorsal ribs are preserved in the blocks, and at least two morphotypes of holocephalous ribs have been identified. The anterior dorsal vertebrae support the larger ribs, which are curved lateroventrally. The capitulum and tuberculum are connected by a narrow flange, forming a single head. In proximal view, the rib attachment twists slightly as an S-shape. The tuberculum is relatively wider than the capitulum, while the middle flange tapers. The rib shaft is curved and has a cylindrical structure with an oblate cross-section. A swollen crest runs along the anterior surface of the shaft, while the posterior surface remains smooth. The last six dorsal vertebrae are associated with smaller ribs, which have slightly curved shafts. Similar to the anterior dorsal ribs, these rib shafts feature a prominent crest on the anterior surface, while the posterior surface is characterized by a longitudinal groove.

## PHYLOGENETIC RESULTS

The current phylogenetic analysis shows significant improvements in diagnosing clades nested within Pareiasauria, especially on the Chinese pareiasaurs. According to the strict consensus tree, several monophyletic clades are identified within the pareiasaurs: the middle Permian Karoo pareiasaurs clade Bradysauria, including *Embrithosaurus*, *Nochelesaurus*, and *Bradysaurus*; the small-sized South African pareiasaurs clade Pumiliopareiasauria, comprinsing *Provelosaurus*, *Nanoparia*, *Pumiliopareia*, and *Anthodon*; the late Permian Pareiasauria group Therischia, which encompasses the horned pareiasaur family Elginiidae such as *Arganaceras*, *Obirkovia*, *Elginia mirabilis*, and *E*. *wuyongae*; Velosauria, a group of derived pareiasaurs as the ancestor of Therischia and Pumiliopareiasauria and all its descendants (Fig. 10).

**FIG. 10.**
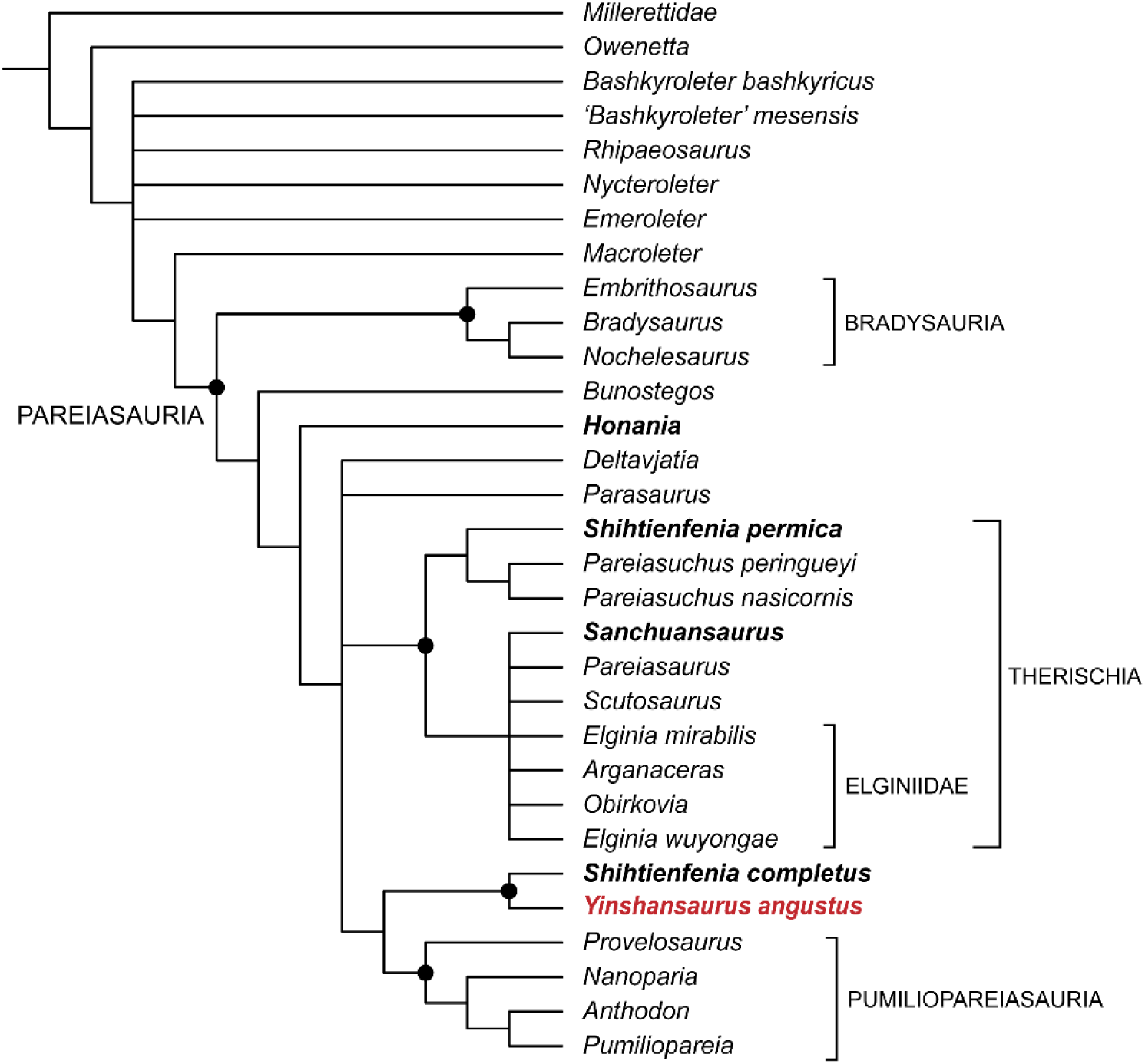
Cladistic relationship of *Yinshansaurus angustus*, strict consensus tree of 210 most parsimonious trees (mpt) with length 297. The Chinese pareiasaurs are bolded. The synapomorphies supportting the main nodes are: Node Therischia: ch49(0 to 1), ch71(0 to 1), ch111(0 to 1), ch124 (1 to 0), ch134(0 to 1), ch136(0 to 2), ch138(0 to 1). Node Pumiliopareiasauria: ch72(1 to 0), ch136(0 to 1). Node of *S*.*completus* and *Y*. *angustus*: ch35 (0 to 1), ch38(1 to 0), ch41(0 to 1), ch54(0 to 1).

The phylogenetic analysis reveals a monophyletic clade that includes *S. completus* and *Y*. *angustus*, which together form the sister group with Pumiliopareiasauria. In contrast, *S*. *permica* remains grouped with *Pareiasuchus*, consistent with previous studies.

In known MPTs, *S. completus* and *Y. angustus* share the following synapomorphies: postfrontal horn, the midline location of the pineal foramen, the exclusion of the postparietal from the margin of the skull, and skull with a of the postorbital length shorter than preorbital length. *Y. angustus* has the following apomorphies: a U-shaped paraoccipital process, a narrower naris dimension, a longer frontal with the length/width ratio of 3.0, vertically oriented maxillary teeth, and 7-9 cusps on the maxillary teeth.

## COMPARISON AND DISSCUSION

### Comparison of Yinshanosaurus with other pareiasaurs

*Yinshanosaurus angustus* shows a combination of morphological characteristics that distinguish it from all other known pareiasaurs. Firstly, the skull displays a high and narrow shape, which is rare in Pareiasauria. The only exception is a narrow skull of *Deltavjatia vjatkensis* (PIN 2212/6) among many wide individuals. However, due to the deviation of its landmarks, Tsuji (2013) suggested that PIN 2212/6 has been laterally compressed. Initially, this type of skull in China was also thought to be the result of compression, but several skulls from different localities suggest this is a natural shape. The high and narrow skull type of *Yinshanosaurus* represents a quite different skull type in pareiasaurs, and it is valid to be named as a new taxon. Consistent with this shape, it has a narrower frontal (with a length/width ratio of 3.0, which is approximately 2.0 in all other pareiasaurs). The U-shaped paraoccipital process also presents in several primitive African pareiasaurs as *Embrithosaurus*, *Nochelesaurus*, *Bradysaurus* (Van den Brandt *et al*. 2023), and *Bunostegos* (Tsuji *et al*. 2013), suggesting this character is irrelevant with the narrow skull shape.

Other than the differentiated characters mentioned above, the distance between the two anterior maxillary foramina is longer in *Y*. *angustus* than that in *S*. *completus* (Fig. 11C, 11D).

**FIG. 11.**
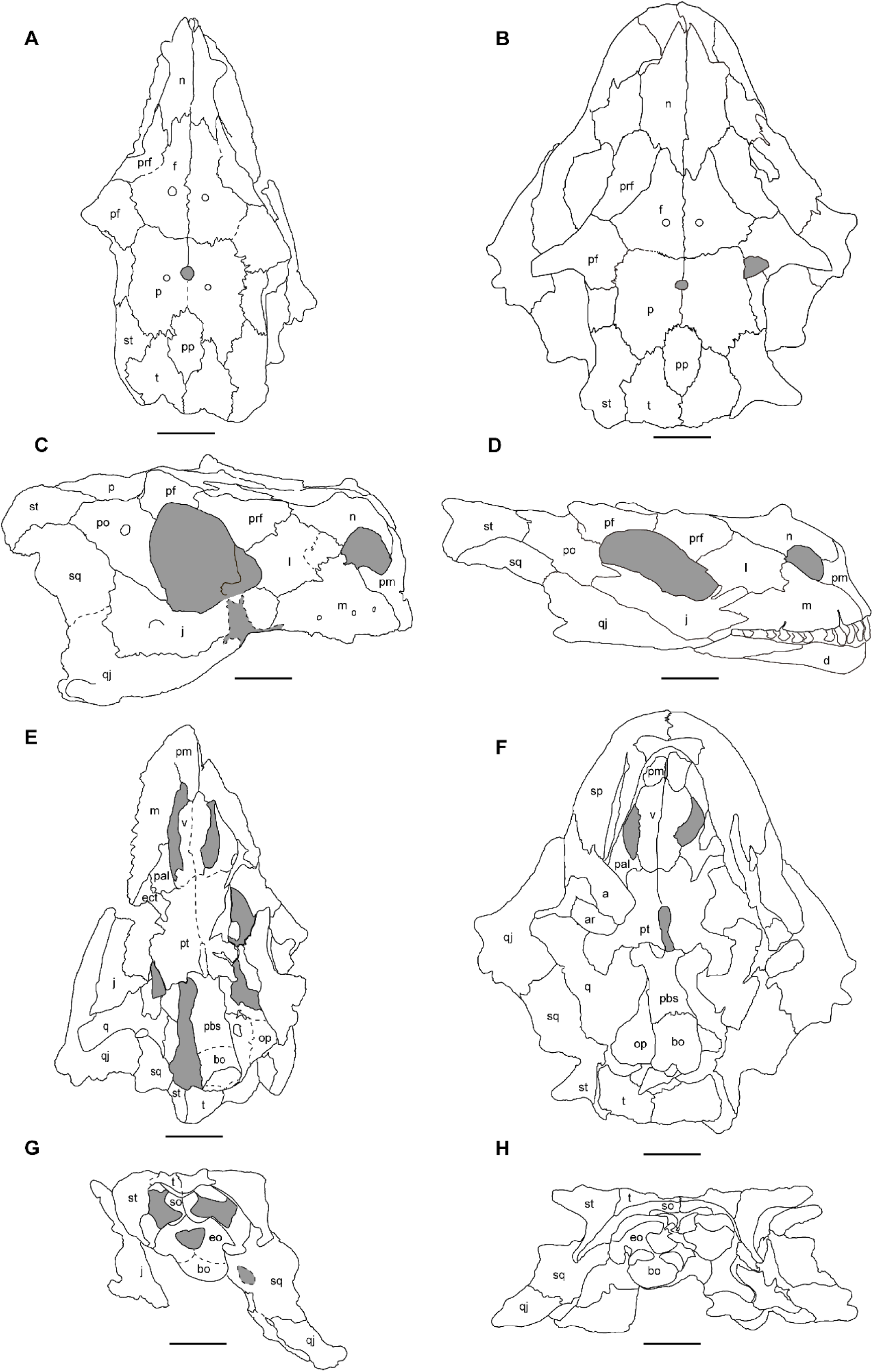
Skull of *Yinshanosaurus angustus* and *Shihtienfenia completus*. in dorsal (A and B), lateral (C and D), ventral (E and F) and occipital (G and H) view. *Abbreviations*: a; angular; ar, articular; bo, bosioccipital; ect; ectopterygoid; eo, exoccipital; f, frontal; j, jugal; l, lacrimal; m, maxilla; n, nasal; op, opisthotic; p, parietal; pal, palatal; pbs, parabasiosphenoid; pf, postfrontal; pm, premaxilla; po, postorbital; pp, postparietal; prf, prefrontal; pt, pterygoid; q, quadrate; qj, quadratojugal; sq, squamosal; so, supraoccipital; st, supratemporal; t, tabular; v, vomer.Scale bars represent 50mm.

**FIG. 12.**
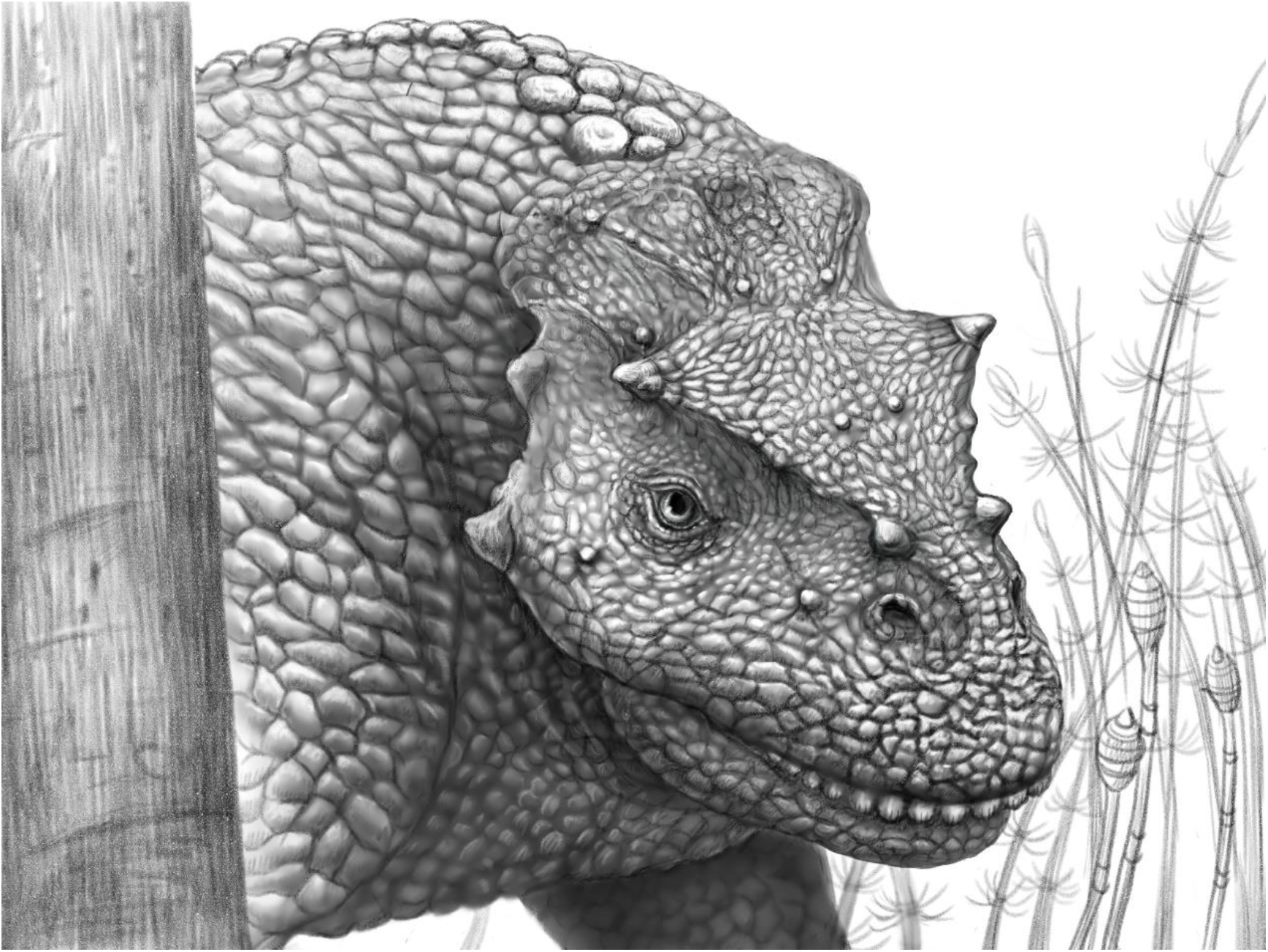
Artists reconstruction of *Yinshanosaurus angustus* sp. nov. (IVPP V33181) in life. Created by Xiao-Cong Guo.

Based on the cranial material, *Yinshanosaurus angustus* cannot be differentiated from *Honania complicidentata*: the maxilla is flat, with vertically oriented maxillary teeth; the quadrate is similar in shape (Xu et al., 2015). Although there are some postcranial bones, they do not provide enough anatomical features. However, they have different positions on current phylogenetic result, and we cannot classify them as same genus. The new species is easily differentiated from *Sanchuansaurus pygmaeus* in which the swelling maxilla bears much more maxillary teeth, and teeth are ventromedially oriented (Gao, 1989). For decades, the derived characteristics of prominent cranial horns and the tabulars suture in midline have been considered exclusive to the Elginiidae. The presence of these features in *S. complete* and *Y*. *angustus* indicates that they are not the autapomorphies of Elginiidae. While the intense compressed parabasisphenoid of *Elginia* (Lee 1997; Liu and Bever 2018) and *Arganaceras* (Jalil and Janvier 2005) distinguish them from *Yinshanosaurus* obviously. As the first pareiasaur in Naobaogou Formation, *Elginia wuyongae*, which represents the dwarf-size pareiasaurs, also features a wide and flat skull shape.

Osteoderms are the most important features of pareiasaurs, these postcranial elements were bony plates of various shapes (Boyarinova and Golubev 2022). The osteoderms were quite uncommon in Chinese pareiasaurs, they are unambiguous present in *Huanghesaurus* (Gao 1983) and *E*. *wuyongae* (Liu and Bever 2018), While *Honania* (Young 1979; Xu *et al*. 2015) and *Shihtienfenia* (Young and Yeh 1963; Wang *et al*. 2019) probably lacks osteoderms. *Y*. *angustus* represents the third osteoderm armored Chinese pareiasaurs, the preserved osteoderms are located along the vertebral column. The outlines of osteoderms are obscure, the overlapped relation is indistinguishable (Fig. 9A, 9C). Therefore, the comparison of osteoderms remains difficult.

The unique shape of the scapular blade distinguishes *Yinshanosaurus* from most other Chinese pareiasaurs. The former exhibits anteroposteriorly aequilatus and strongly torsional scapulars, while the latter display a noticeable distal expansion on the straight scapular blade (flared).

### The validity of Shihtienfenia completus

The first complete Chinese pareiasaur skull of *Shihtienfenia completus* was briefly described in Chinese (Wang *et al*. 2019). Based on the postparietal is excluded from the margin of the skull table, *S. completus* was tentatively classified in Elginiidae. In this study, *S*. *completus* is recovered as a species not closely related to *S*. *permica*. If this is the real relationship between them, the generic name of *Shihtienfenia completus* need to be changed. So, we do not refer the new species to *Shihtienfenia*.

The relationship between *S*. *permica* and *S. completus* may be resolved later after the detailed study on the abundant disarticulated postcranial elements (SXNHM V0010) excavated together with the only known specimen of *S. completus*.

### The age estimation and geography distribution of Yinshanosaurus

Although Pareiasaurs existed from the Guadalupian to Lopingian, only Bradysauria thrived during the mid-Permian (Van den Brandt *et al*. 2022). Most Chinese pareiasaurs are produced from the Upper Shihhotse and Sunjiagou formations in the Ordos Basin (Li *et al*. 2008; Liu 2018). Although new dating of tuff from the marker sandstone layer in the Upper Shihhotse Formation constrains the tetrapod fossils age from later than 261.75Ma in North China (Wu *et al*. 2021), differences of paleolatitude and tetrapods fauna suggest limited comparability between the Sunjiagou and Naobaogou Formations.

The age of the Naobaogou Formation is generally regarded as late Permian based on vertebrate biostratigraphy. Member I of the Naobaogou Formation may be classified as Wuchiapingian (259.1-254.1Ma) based on the presence of *Euchambersia* (Liu and Abdala 2022). Given that *Y*. *angustus* is the sister species of *S. completus*, their age should be concurrent. Therefore, the age of the Sunjiagou Formation is supposed to be Wuchiapingian rather than Changhsingian.

## CONCLUSION

*Yinshanosaurus angustus* is validated as a new taxon based on a combination of characters: skull high and narrow, with the length of the skull table exceeding the width between the two quadratojugals, long frontal with a length-to-width ratio of ∼3.0, U-shaped paraoccipital process in occipital view, maxillary teeth oriented vertically, with only 7-9 cusps. The skeleton of *Y*. *angustus* provides the complete cranial and articulated postcranial details of Chinese pareiasaurs for the first time.

In phylogeny, *Y*. *angustus* and *S*. *completus* form a monophyletic clade characterized by a unique combination of features, including the presence of a prominent postfrontal horn, a pineal foramen located midway along the interparietal suture, and tabular sutures in the midline that encircle the postparietal. Furthermore, the postorbital region of the skull is shorter than the preorbital region. This newly identified monophyletic group forms a sister clade to Pumiliopareiasauria, distinguishing it from other pareiasaurs.

Therefore, three preliminary clades of Chinese pareiasaurs have been established to date: *Honania*, which is found in the lowest stratigraphic horizon and occupies a basal position in the consensus tree; *S*. *permica* and *Sanchuansaurus*, both of which belong to Therischia; along with the new monophyly consist of *S*. *completus* and *Y*. *angustus*. Although the newly discovered complete skull has enhanced the framework, the controversy surrounding the phylogenetic relationships of Chinese pareiasaurs, which is attributed to the indirectly compared elements still need improvement. Further studies are necessary to confirm the stable positions of the remaining Chinese species based on postcranial elements of *S*. *completus*.

## Acknowledgements.

We gratefully acknowledge the hard work of the 2018 field team (L Li, Y-D Liu, Y-F Liu, P-J Zhang and S-G Zhang), Y-D Liu and H-L Fu prepared the specimens. W Gao took the photos. This work was supported by the Strategic Priority Research Program of Chinese Academy of Sciences (XDB26000000).

## Author contributions

Conceptualization Jian Yi (JY), Jun Liu (JL); Data Curation JY; Formal Analysis JY; Investigation JY, JL; Methodology JL; Visualization JY; Writing – Original Draft Preparation YJ; Writing – Review & Editing JL.

## Appendix 1 Data Matrix used in phylogenetic analysis. MATRIX

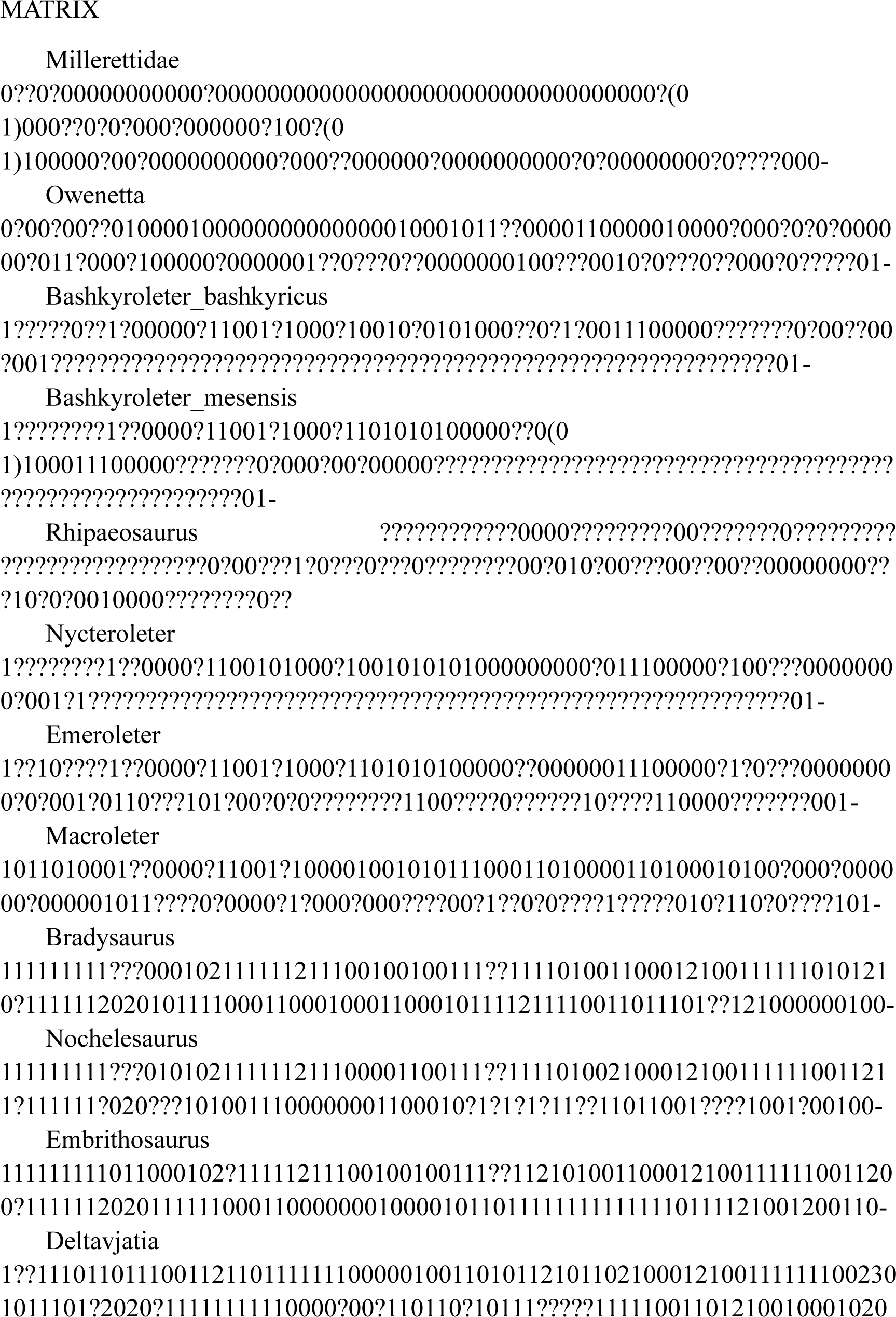
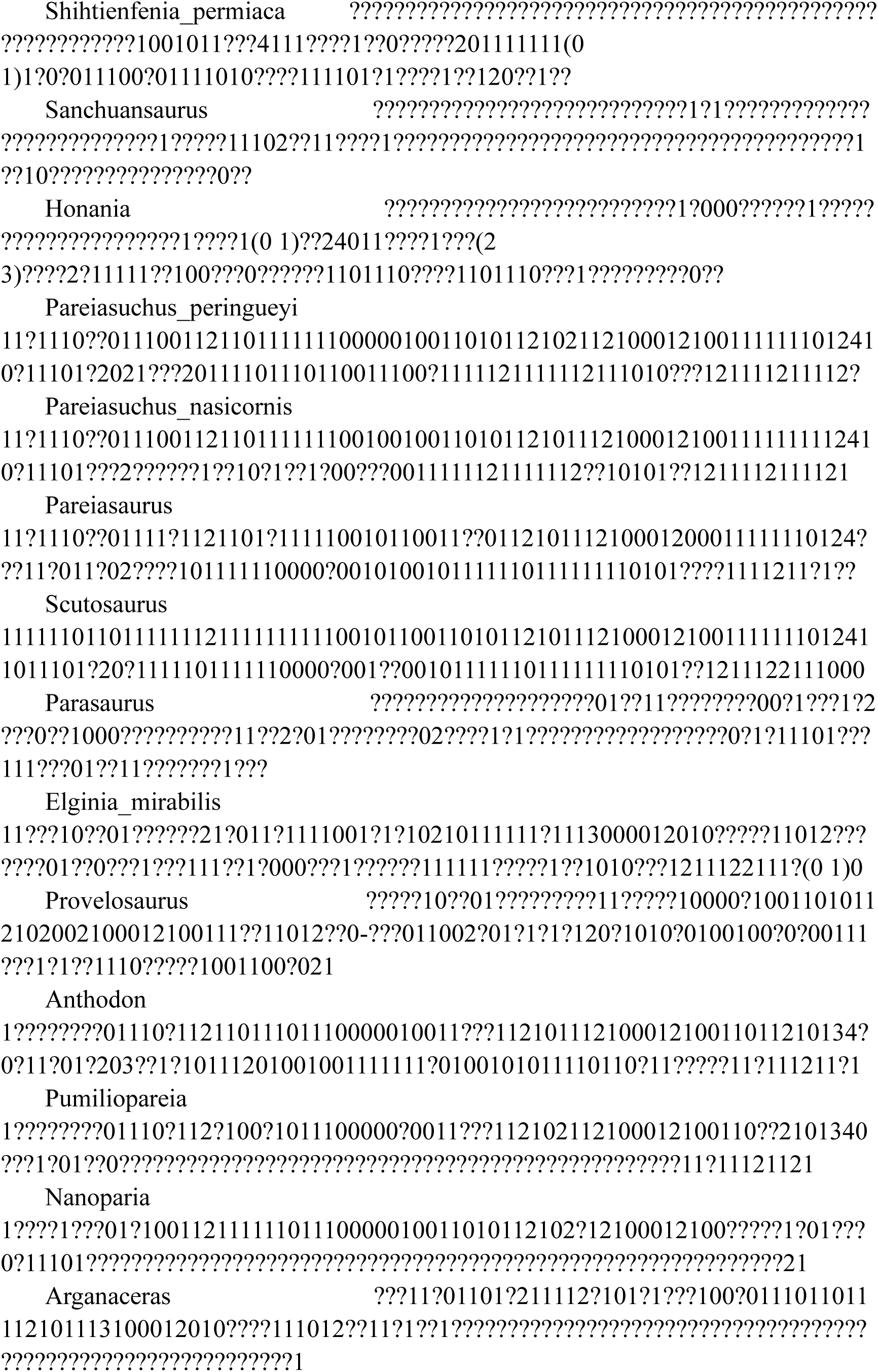
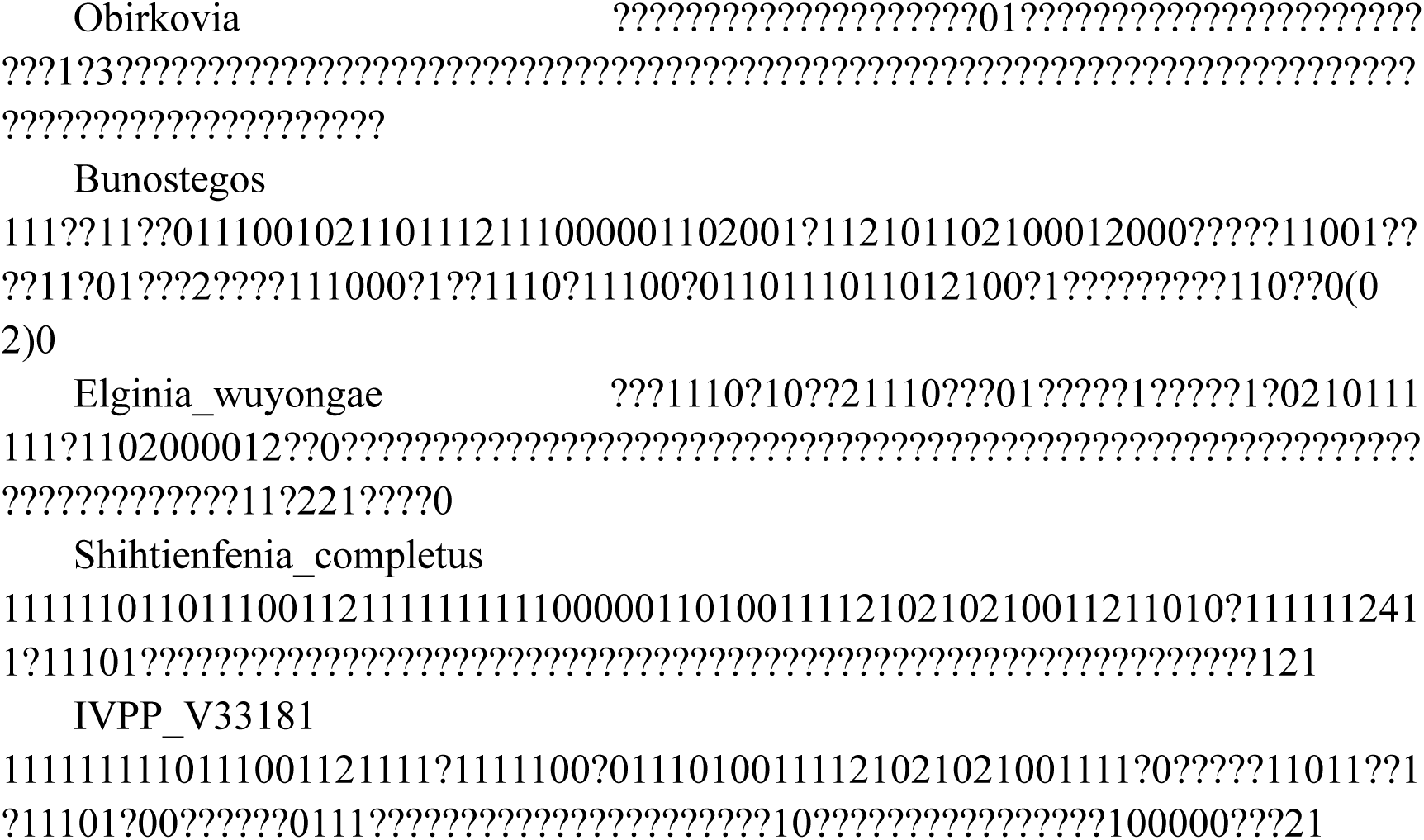

## Appendix 2 SUPPLEMENTAL MATERIAL: CHARACTER LIST

List of characters used in the phylogenetic analysis.

We used the character list of Van den Brandt *et al.,* (2023) and modified it by; re-wording **characters 54**: “Postorbital region of skull, length: length at least equals preorbital region of the skull (0); postorbital region shorter than preorbital region (1)”. and by introducing an additional character state (state 2) for **character 56**, “Frontal shape: frontal length reduced, three times as long as wide (1)” for *Yinshanosaurus*,and revised the state “frontals short, with a length not more than two times the width” from 1 to 2.

**Braincase (characters 1-28, except 21, 22)**

1. Basicranial articulation: Pterygoid and/or epipterygoid articulation with the basipterygoid process is mobile (0); articulation is immobile (1). (Jalil & Janvier, 2005:1; Tsuji, 2013:1; Turner *et al.,* 2015:1; Van den Brandt *et al.,* 2023:1)

2. Sphenethmoid, ossification: absent (0); present (1). (Jalil & Janvier, 2005:2; Tsuji, 2013:2; Turner *et al.,* 2015:2; Van den Brandt *et al.,* 2023:2)

3. Prootic, medial wall, ossification: absent (0): present (1). (Jalil & Janvier, 2005:3; Tsuji, 2013:3; Turner *et al.,* 2015:3; Van den Brandt *et al.,* 2023:3)

4. Exoccipital, lateral flange: absent (0); present (1). (Jalil & Janvier, 2005:4; Tsuji, 2013:4; Turner *et al.,* 2015:4; Van den Brandt *et al.,* 2023:4)

5. Exoccipital, lateral flange, size: small flange (0); lateral flange of the exoccipital well-developed and extends well along the paroccipital process of the opisthotic (1). (Jalil & Janvier, 2005:4; Tsuji, 2013:4; Turner *et al.,* 2015:5; Van den Brandt *et al.,* 2023:5)

6. Paroccipital process, suture: not sutured to the squamosal and supratemporal (0); paroccipital process of the opisthotic is antero-posteriorly expanded and sutured to ventrally-directed flange from the squamosal and supratemporal (1). (Jalil & Janvier, 2005:5; Tsuji, 2013:5; Turner *et al.,* 2015:6; Van den Brandt *et al.,* 2023:6)

7. Paroccipital process, orientation: projects laterally from the neurocranium (0); is U-shaped in occipital view (1). (Jalil & Janvier, 2005:6; Tsuji, 2013:6; Turner *et al.,* 2015:7; Van den Brandt *et al.,* 2023:7)

8. Ventral otic fissure: present (0); absent (1). (Jalil & Janvier, 2005:7; Tsuji, 2013:7; Turner *et al.,* 2015:8; Van den Brandt *et al.,* 2023:8)

9. Parabasisphenoid and basioccipital, braincase floor: not thickened (0); thickened (1). (Jalil & Janvier, 2005:8; Tsuji, 2013:8; Turner *et al.,* 2015:9; Van den Brandt *et al.,* 2023:9)

10. Cultriform process: present (0); absent (1). (Jalil & Janvier, 2005:9; Tsuji, 2013:9; Turner *et al.,* 2015:10; Van den Brandt *et al.,* 2023:10)

11. Cultriform process, length: relatively long, more than the half of the distance between the anterior extent of the rostrum (tip of snout) to the basal tubera (0); short, less than a third of this distance (1). (Jalil & Janvier, 2005:9; Tsuji, 2013:9; Turner *et al.,* 2015:11; Van den Brandt *et al.,* 2023:11)

12. Cultriform process, shape of anterior tip: pointed (0); blunt (1). (Jalil & Janvier, 2005:10; Tsuji, 2013:10; Turner *et al.,* 2015:12; Van den Brandt *et al.,* 2023:12)

13. Basisphenoid, body: wide, strongly constricted, giving it an hourglass shape in ventral view (0); wide, not strongly laterally constricted (1); narrow with relatively straight margin (2). (Jalil & Janvier, 2005:11; Tsuji, 2013:11; Turner *et al.,* 2015:13; Liu & Bever, 2018:13 modified; Van den Brandt *et al.,* 2023:13)

14. Basisphenoid, ventral surface of each basipterygoid process, turbercles: featureless, lacking tubercles (0); tubercles present on the ventral surface of each basipterygoid process (basisphenoid), immediately posterior to the interpterygoid vacuity and the transverse suture with the pterygoid (1). (Jalil & Janvier, 2005:12; Tsuji, 2013:12; Turner *et al.,* 2015:14; Van den Brandt *et al.,* 2022:14 modified; Van den Brandt *et al.,* 2023:14)

15. Basioccipital, ventral surface, central boss: absent (0); present (1). (Jalil & Janvier, 2005:13; Tsuji, 2013:13; Turner *et al.,* 2015:15; Van den Brandt *et al.,* 2023:15)

16. Basal tubera: absent (0); present (1). (Jalil & Janvier, 2005:14; Tsuji, 2013:14; Turner *et al.,* 2015:16; Van den Brandt *et al.,* 2023:16)

17. Basal tubera, position: situated posteriorly, closer to the occipital condyle than the basipterygoid process (0); basal tubercles situated approximately midway between the occipital condyle and the basipterygoid processes, or even further anteriorly (1). (Jalil & Janvier, 2005:15; Tsuji, 2013:15; Turner *et al.,* 2015:17; Van den Brandt *et al.,* 2023:17)

18. Choana, shape: situated in a lateral position, bounded laterally by the maxilla, diverge posteriorly, parallel to tooth row throughout (0); parallel, positioned more medially, delimited posterolaterally by the palatine (1); choanae even more medially positioned, with the palatine constituting more than 50% of the lateral border, medial border formed entirely by the vomer (2). (Jalil & Janvier, 2005:16; Tsuji, 2013:16; Turner *et al.,* 2015:18; Van den Brandt *et al.,* 2023:18)

19. Vomer, alar flange (lateral flange): absent (0); present (1). (Jalil & Janvier, 2005:17; Tsuji, 2013:17; Turner *et al.,* 2015:19; Van den Brandt *et al.,* 2023:19)

20. Foramen palatinum posterius, size: small or absent and delineated by the bones of the skull roof (0); large, medially positioned and defined by the palatine and the ectopterygoid without participation of the bones of the skull roof (1). (Jalil & Janvier, 2005:18; Tsuji, 2013:18; Turner *et al.,* 2015:20; Van den Brandt *et al.,* 2023:20)

21. Skull roof, radiating ridges: dermal sculpturing in the form of relatively straight ridges radiating from the center of dermal skull roof bones absent (0); regular ridges present (1). (Tsuji, 2013:115; Turner *et al.,* 2015:21; Van den Brandt *et al.,* 2023:21)

22. Circular pits: cranial sculpture in the form of circular pits absent (0); present (1). (Tsuji, 2013:116; Turner *et al.,* 2015:22; Van den Brandt *et al.,* 2023:22)

23. Medial prepalatal foramen bordered by the premaxilla and the vomer: absent (0); present (1). (Jalil & Janvier, 2005:19; Tsuji, 2013:19; Turner *et al.,* 2015:23; Van den Brandt *et al.,* 2023:23)

24. Interpterygoid vacuity, length: long, at least 15% of skull length (0) short, less than 15% of skull length (1). (Tsuji, 2013:20; Turner *et al.,* 2015:24; Van den Brandt *et al.,* 2023:24)

25. Interpterygoid vacuity, anterior shape: V-shaped, extends far anteriorly and is anteriorly pointed (0); anterior border is upside-down U-shaped, anteriorly bulged (pointing) or convex (1); transversely-oriented, anterior border is U-shaped or posteriorly bulged or convex (2).(Jalil & Janvier, 2005:20; Tsuji, 2013:21; Turner *et al.,* 2015:25; Van den Brandt *et al.,* 2023:25 modified; Van den Brandt *et al.,* 2023:25)

26. Pterygoid, transverse flange, shape: large and directed laterally (0); reduced, directed more anteriorly than laterally, without contact with the cheek (1). (Jalil & Janvier, 2005:21; Tsuji, 2013:22; Turner *et al.,* 2015:26; Van den Brandt *et al.,* 2023:26)

27. Pterygoid, transverse flange, orientation: extends ventrally below the level of the alveolar ridge (0); oriented primarily horizontally, so the level of the palate is higher, not reaching the level of the alveolar ridge (1). (Jalil & Janvier, 2005:22; Tsuji, 2013:23; Turner *et al.,* 2015:27; Van den Brandt *et al.,* 2023:27)

28. Supraoccipital, form: large, with longitudinal contact with the postparietal (0); high and narrow, forming along all of its length a solid sagittal suture with a ventral projection of the postparietal (1). (Jalil & Janvier, 2005:23; Tsuji, 2013:24; Turner *et al.,* 2015:28; Van den Brandt *et al.,* 2023:28)

**Skull Roof (characters 29-58, and 21, 22)**

29. External naris, form: round and small (0); very anteroposteriorly elongate (1). (Jalil & Janvier, 2005:25; Tsuji, 2013:25; Turner *et al.,* 2015:29; Van den Brandt *et al.,* 2023:29)

30. External naris, maxilla contribution: maxilla either excluded from naris or forms only its ventral/posterior edge (0); maxilla extends also to the posterodorsal margin of naris (1). (Tsuji, 2013:125; Turner *et al.,* 2015:30; Van den Brandt *et al.,* 2023:30)

31. Maxilla, boss: a boss or horn on the maxilla immediately posterior to the external naris feebly developed or absent (0); prominent boss or horn present (1). (Jalil & Janvier, 2005:27; Tsuji, 2013:26; Turner *et al.,* 2015:31, Van den Brandt *et al.,* 2022:31 modified; Van den Brandt *et al.,* 2023:31)

32. Maxilla, dorsal process: reduced, not reaching the nasal, so that the lacrimal contributes to the ventral border of the naris (0); large anterodorsal extension of the maxilla, excluding the lacrimal from the external naris (1). (Jalil & Janvier, 2005:28; Tsuji, 2013:27; Turner *et al.,* 2015:32; Van den Brandt *et al.,* 2023:32 modified; Van den Brandt *et al.,* 2023:32)

33. Snout dimensions (anteriorly): broader than high (0); as high as wide (1). (Lee, 1997:30; Jalil & Janvier, 2005:31; Tsuji, 2013:28; Turner *et al.,* 2015:33)

34. Postfrontal, shape: mediolaterally narrow, more than 2 times as long as wide, contributes to the orbital margin (0); widened mediolaterally, around 2 times as wide as long, no or only feeble contribution to the orbital rim (1). (Jalil & Janvier, 2005:32; Tsuji, 2013:29; Turner *et al.,* 2015:34 modified; Van den Brandt *et al.,* 2023:34)

35. Postfrontal, ‘horn’: absent (0); present (1). (Tsuji *et al.,* 2013:126; Turner *et al.,* 2015:35; Van den Brandt *et al.,* 2023:35)

36. Orbit, shape: circular, no posterior emargination (0); posterior emargination of orbits (1). (Lee, 1995:23, Lee, 1997:32; Jalil & Janvier, 2005:33; Tsuji, 2013:30; Turner *et al.,* 2015:36; Van den Brandt *et al.,* 2023:36)

37. Circumorbital tuberocities: circumorbital skull elements lacking tubercles or bosses (0); circumorbital tubercles small (1); circumorbital tubercles large (2). (Tsuji, 2013:114; Turner *et al.,* 2015:37, Liu & Bever, 2018:37 modified; Van den Brandt *et al.,* 2023:37)

38. Pineal foramen, position: pineal foramen situated about halfway along the interparietal suture (0); placed more anteriorly, close to the frontal-parietal suture (1). (Jalil & Janvier, 2005:34 & 35; Tsuji, 2013:31; Turner *et al.,* 2015:38; Van den Brandt *et al.,* 2023:38)

39. Tabular (supernumerary element): present (0); absent (1). (Jalil & Janvier, 2005:37; Tsuji, 2013:32; Turner *et al.,* 2015:39; Van den Brandt *et al.,* 2023:39)

40. Tabular (supernumerary element), form: small, largely an occipital element (0); integrated into skull table (1). (Jalil & Janvier, 2005:37; Tsuji, 2013:32; Turner *et al.,* 2015:40; Van den Brandt *et al.,* 2023:40)

41. Tabular (supernumerary element), contact: do not contact each other posteriorly (0); very well-developed, make contact posteriorly, excluding the postparietals from the posterior edge of the skull table (1). (Modified from Jalil & Janvier, 2005:38; Tsuji, 2013:33; Turner *et al.,* 2015:41; Van den Brandt *et al.,* 2023:41)

42. Postparietal, form: small, largely an occipital element (0); integrated into skull table (1). (Jalil & Janvier, 2005:39; Tsuji, 2013:34; Turner *et al.,* 2015:42; Van den Brandt *et al.,* 2023:42)

43. Postparietal, fusion: paired (0); fused into a single element and exposed well dorsally (1). (Jalil & Janvier, 2005:40; Tsuji, 2013:35; Turner *et al.,* 2015:43; Van den Brandt *et al.,* 2023:43)

44. Expanded quadratojugal (cheek) flange: absent, no cheek flange, quadratojugal (cheek) does not extend below level of tooth row, the ventral surface of the quadratojugal is continuous with and forms a straight edge with that of the maxilla (0); small cheek flange present, quadratojugal (cheek) flange extends below the level of the maxillary tooth row 0° – 40° (1); large cheek flange present, quadratojugal (cheek) flange extends below the level of the maxillary tooth row 41° or more (2). (Jalil & Janvier, 2005:46; Tsuji, 2013:38; Turner *et al.,* 2015:44; Liu & Bever, 2018:44; Van den Brandt *et al.,* 2020, 2023:44 modified; Van den Brandt *et al.,* 2023:44)

45. Jugal, anterior process: does not extend to anterior orbital rim (0); extends at least to level of orbital rim (1). (Tsuji, 2013:117; Turner *et al.,* 2015:45; Van den Brandt *et al.,* 2023:45)

46. Jugal, suture with maxilla: straight, jugal thins out smoothly towards anterior, or upright, but no dramatic vertical ’step’ (0); “stepped”, anteriormost tip of jugal very narrow but expands broadly posteriorly along with a dramatic thinning of the posterior process of the maxilla (1). (Tsuji, 2013:123; Turner *et al.,* 2015:46; Van den Brandt *et al.,* 2023:46)

47. Quadratojugal, anterior extent: does not reach level of posterior border of orbit (0); reaches posterior border of orbit (1); reaches or almost reaches the anterior margin of the orbit (2). (Tsuji, 2013:118; Turner *et al.,* 2015:47; Van den Brandt *et al.,* 2023:47 modified; Van den Brandt *et al.,* 2023:47)

48. Quadratojugal, ventral margin, ornamentation (bosses): absent, no ornamentation (bosses) on the ventral margin of the quadratojugal (0); present, ornamentation (bosses) on the ventral margin of the quadratojugal (1). (Jalil & Janvier, 2005:48; Tsuji, 2013:40; Turner *et al.,* 2015:48; Van den Brandt *et al.,* 2020:48 modified; Van den Brandt *et al.,* 2023:48)

49. Junction of skull table and cheek: both flat surfaces, form a distinct angle where they meet, particularly posterior to the orbits (0); postorbital portion of this junction is rounded, no clear edge between these two surfaces posterior to the orbit (1). (Jalil & Janvier, 2005:43; Tsuji, 2013:36; Turner *et al.,* 2015:49, Van den Brandt *et al.,* 2022:49 modified; Van den Brandt *et al.,* 2023:49)

50. Cheek ornamentation style (quadratojugal and squamosal): no ornamentation on the posterior or ventral margins of the cheek (0); ornamentation present on the posterior or ventral margins of the cheek in the form of low rounded bosses (1); ornamentation present on the posterior or ventral margins of the cheek in the form of well-developed, more distinct, taller, more pointed bosses (2); ornamentation present on the posterior or ventral margins of the cheek in the form of prominent, conical horns with sharp, pointed tips (3). (Jalil & Janvier, 2005:47; Tsuji, 2013:39; Turner *et al.,* 2015:50, Van den Brandt *et al.,* 2020:50 modified; Van den Brandt *et al.,* 2023:50)

51. Temporal emargination (otic notch): absent, or very small (0), emargination present on the posterior border of the cheek, in the dorsal portion of the squamosal, just below the occipital shelf of the supratemporal, being horizontal and extending medially along the internal surface of the squamosal (1). (Jalil & Janvier, 2005:45; Tsuji, 2013:37; Turner *et al.,* 2015:51, Liu & Bever, 2018:51; Van den Brandt *et al.,* 2020:51 modified51; Van den Brandt *et al.,* 2023:)

52. Temporal, otic notch, position: restricted to posterior half of cheek (0); closely approaches the orbital margin (1). (Tsuji, 2013:124; Turner *et al.,* 2015:52; Van den Brandt *et al.,* 2023:52)

53. Temporal, ventral emargination: absent (0); present (1). (Tsuji, 2013:41; Turner *et al*.; Van den Brandt *et al.,* 2023:53)

54. Postorbital region of skull, length: length at least equals preorbital region of the skull (0); postorbital region shorter than preorbital region (1) **(modified).** (Modified from Laurin & Reisz, 1995:32; Tsuji, 2013:42; Turner *et al.,* 2015:54; Van den Brandt *et al.,* 2023:54)

55. Frontal, contribution to the orbit: present (0); frontals excluded from the orbit by contact between the prefrontal and postfrontal (1). (Jalil & Janvier, 2005:49; Tsuji, 2013:43; Turner *et al.,* 2015:55; Van den Brandt *et al.,* 2023:55)

56. Frontal, shape: slim and long, four times as long as wide (0); frontal length reduced, three times as long as wide (1); frontals short, with a length not more than two times the width (2) **(modified).** (Jalil & Janvier, 2005:50; Tsuji, 2013:44; Turner *et al.,* 2015:56; Van den Brandt *et al.,* 2023:56)

57. Frontal, central boss: absent (0); present (1). (Tsuji *et al.,* 2013:127; Turner *et al.,* 2015:57; Van den Brandt *et al.,* 2023:57)

58. Boss ornamentation: dermal bosses of skull bones have no central pointed horn (0); dermal bosses of cranial bones have a central long, pointed horn (1). (Jalil & Janvier, 2005:51; Tsuji, 2013:45; Turner *et al.,* 2015:58; Van den Brandt *et al.,* 2023:58)

**Lower jaw (characters 59-64)**

59. Jaw articulation, position: anterior to occiput (0); even with occiput (1). (Tsuji, 2013:120; Turner *et al.,* 2015:59; Van den Brandt *et al.,* 2023:59)

60. Mandibular symphysis: splenial is excluded from the mandibular symphysis (0); splenial forms the ventral portion of the mandibular symphysis (1). (Jalil & Janvier, 2005:52; Tsuji, 2013:46; Turner *et al.,* 2015:60; Van den Brandt *et al.,* 2023:60)

61. Angular, boss: absent, ventral surface of the angular is smooth (0); boss is present (1). (Jalil & Janvier, 2005:53; Tsuji, 2013:47; Turner *et al.,* 2015:61; Van den Brandt *et al.,* 2023:61)

62. Angular, boss, form: low and rounded (0, prev. 1); well developed with a prominent, pointed tubercle (1, prev. 2). (Jalil & Janvier, 2005:53; Tsuji, 2013:47; Turner *et al.,* 2015:62; Van den Brandt *et al.,* 2023:62)

63. Articular, retroarticular process, dorsal projection: without a projection, tapers gradually to end (0); small projection (“dorsal lump” of Lee, 1997) present at the very posterior end of the retroarticular process (1). (Jalil & Janvier, 2005:55; Tsuji, 2013:48; Turner *et al.,* 2015:63; Van den Brandt *et al.,* 2023:63)

64. Articular (region), lateral shelf: absent, the lateral surface of the articular region is smooth (0); present, there is a lateral extension of the surangular or articular, the lateral surface of the effected element extends dorsolaterally (1). (Modified slightly from Jalil & Janvier, 2005:56; Tsuji, 2013:49; Turner *et al.,* 2015:64; Van den Brandt *et al.,* 2023:64)

**Dentition (characters 65-73)**

65. Teeth, labiolingual compression (anteroposteriorly expanded): teeth not labio-lingually compressed (0); teeth labio-lingually compressed, leaf-shaped, with small denticles on the tooth crown (1); labio-lingual compression very pronounced, giving the marginal teeth a fan shape (2). (Jalil & Janvier, 2005:60; Tsuji, 2013:52; Turner *et al.,* 2015:65; Van den Brandt *et al.,* 2023:65)

66. Teeth, cusp arrangement: three central cusps close together, more lateral cusps farther apart and spaced farther apart from each other than these central three (0); cusps regularly spaced along the tooth crown (1). (Jalil & Janvier, 2005:61; Tsuji, 2013:53; Turner *et al.,* 2015:66; Van den Brandt *et al.,* 2023:66)

67. Maxillary teeth, orientation: maxillary teeth oriented vertically, teeth point directly downwards (0); alveolar ridge inflected towards the palate, teeth oriented ventromedially

(1). (Jalil & Janvier, 2005:57; Tsuji, 2013:50; Turner *et al.,* 2015:67; Van den Brandt *et al.,* 2023:67)

68. Upper jaw teeth (premaxilla and maxilla), number on each side: 20 or more (0), 18 or less

(1). (Xu *et al.,* 2015:51; Turner *et al.,* 2015:68; Van den Brandt *et al.,* 2020:68 modified; Van den Brandt *et al.,* 2023:68)

69. Maxillary teeth, cusp number: conical, single cusp (0); 7–9 cusps on each maxillary tooth (1); 9–11 cusps (2); more than 11 cusps (3). (Jalil & Janvier, 2005:62; Tsuji, 2013:54; Turner *et al.,* 2015:69; Van den Brandt *et al.,* 2022:69 modified; Van den Brandt *et al.,* 2023:69)

70. Mandibular teeth, cusp number: conical, without cusps (0); 2–7 cusps on each mandibular tooth (1); 7–9 cusps (2); 9–11 cusps (3); more than 11 cusps (4). (Jalil & Janvier, 2005:63; Tsuji, 2013:55; Turner *et al.,* 2015:70; Van den Brandt *et al.,* 2023:70)

71. Mandibular teeth, lingual surface, shape: smooth (0); has a distinct, triangular lingual ridge, narrowing towards the crown of the tooth (1). (Jalil & Janvier, 2005:64; Tsuji, 2013:56; Turner *et al.,* 2015:71; Van den Brandt *et al.,* 2023:71)

72. Marginal teeth, lingual surface, horizontal cingulum: absent (0); present (1). (Xu *et al.,* 2015:57; Turner *et al.,* 2015:72; Van den Brandt *et al.,* 2023:72)

73. Marginal teeth, lingual surface, cingulum, extent: on some maxillary teeth (0); on most maxillary teeth (1). (Xu *et al.,* 2015:57; Turner *et al.,* 2015:73; Van den Brandt *et al.,* 2023:73)

**Palate (characters 74-78)**

74. Pterygoid, transverse flange, dentition: no dentition on the transverse flange of the pterygoid (0); teeth present on the transverse flange of the pterygoid (1). (Jalil & Janvier, 2005:66; Tsuji, 2013:58; Turner *et al.,* 2015:74; Van den Brandt *et al.,* 2022:74 modified; Van den Brandt *et al.,* 2023:74)

75. Pterygoid, anterior extent: reaches level of choana (0); posterior to choana (1). (Tsuji, 2013:113; Turner *et al.,* 2015:75; Van den Brandt *et al.,* 2023:75)

76. Pterygoid, quadrate ramus: merges smoothly into transverse flange without distinctive excavation (0); deep excavation on posterolateral surface (1). (Tsuji, 2013:122; Turner *et al.,* 2015:76; Van den Brandt *et al.,* 2023:76)

77. Palatal teeth: medial rows of palatal denticles parallel and close together and to the medial axis of the skull (0); medial rows of palatal denticles widely separated, converging anteriorly (1). (Jalil & Janvier, 2005:67; Tsuji, 2013:59; Turner *et al.,* 2015:77; Van den Brandt *et al.,* 2023:77)

78. Caniniform region: present (0); absent (1). (Tsuji, 2013:119; Turner *et al.,* 2015:78; Van den Brandt *et al.,* 2023:78)

**Vertebrae (characters 79-86)**

79. Atlas-axis fusion: pleurocentrum of the atlas and axial intercentrum fused (0); atlas pleurocentrum separate from the axial intercentrum (1). (Jalil & Janvier, 2005:69; Tsuji, 2013:61; Turner *et al.,* 2015:79; Cisneros *et al.,* 2021: 79 polarity reversed; Van den Brandt *et al.,* 2023:79)

80. Presacral vertebrae, number: more than 20 presacral vertebrae (0); 20 presacral vertebrae (1); 19 or fewer presacral vertebrae (2). (Jalil & Janvier, 2005:68; Tsuji, 2013:60; Turner *et al.,* 2015:80; Van den Brandt *et al.,* 2023:80)

81. Lumbar vertebrae: absent (0); present (1). (Jalil & Janvier, 2005:70; Tsuji, 2013:62; Turner *et al.,* 2015:81; Van den Brandt *et al.,* 2023:81)

82. Sacral vertebrae, number: two (0); three (1); four (2); five (3). (Jalil & Janvier, 2005:71; Tsuji, 2013:63; Turner *et al.,* 2015:82; Van den Brandt *et al.,* 2023:82)

83. Caudal vertebrae, number (tail length): long, with more than 25 caudal vertebrae (0); short, less than 25 caudal vertebrae (1). (Jalil & Janvier, 2005:73; Tsuji, 2013:64; Turner *et al.,* 2015:83; Van den Brandt *et al.,* 2023:83)

84. Caudal vertebrae, lateral projections: generally present on the first five but never on more than 9 of the first (most anterior) caudal vertebrae (0); prominent lateral projections on at least the first 9 caudal vertebrae (1). (Jalil & Janvier, 2005:74; Tsuji, 2013:65; Turner *et al.,* 2015:84; Van den Brandt *et al.,* 2023:84)

85. Caudal vertebrae, lateral projections, shape: projections form an ’L’, as their distal portions are recurved posteriorly parallel to the axis of the body (0); projections almost straight and directed laterally (1). (Jalil & Janvier, 2005:75; Tsuji, 2013:66; Turner *et al.,* 2015:85; Van den Brandt *et al.,* 2023:85)

86. Caudal vertebrae, hemal arch, insertion: between two caudal vertebrae (0); articulate with only one centrum via a facet of articulation found on posteroventral projections of the centra (1). (Jalil & Janvier, 2005:76; Tsuji, 2013:67; Turner *et al.,* 2015:86; Van den Brandt *et al.,* 2023:86)

**Scapulocoracoid (characters 87-91)**

87. Scapula, blade, length: less than two times the glenoid fossa diameter (0), between two and three times the glenoid fossa diameter (1), greater than or equal to three times the glenoid fossa diameter (2). (Lee, 1993:B2, Lee, 1995:41, Lee, 1997:75; Laurin & Reisz, 1995:96; Jalil & Janvier, 2005:78; Tsuji, 2013:69; Turner *et al.,* 2015:87)

88. Scapula, blade, shape: straight preaxial and postaxial margins expanding gradually (0); preaxial and postaxial margins are curved, expansion pronounced dorsally at the distal end (flared) (1). (Turner *et al.,* 2015:88; Van den Brandt *et al.,* 2023:88)

89. Acromion process: absent (0), present (1). (Jalil & Janvier, 2005:77; Tsuji, 2013:68; Turner *et al.,* 2015:89; Van den Brandt *et al.,* 2023:89)

90. Posterior coracoid, dorsal edge: almost horizontal, meets the posterior border of the scapula at an angle of less than 135° (0); dorsal edge of the posterior coracoid is oriented posteroventrally, forms an angle of more than 135° with the posterior border of the scapula (1). (Jalil & Janvier, 2005:80; Tsuji, 2013:70; Turner *et al.,* 2015:90; Van den Brandt *et al.,* 2023:90)

91. Cleithrum: present (0); absent (1). (Jalil & Janvier, 2005:81; Tsuji, 2013:71; Turner *et al.,* 2015:91; Van den Brandt *et al.,* 2023:91)

**Humerus (characters 92-104)**

92. Humerus, torsion: the planes of proximal and distal expansion makes an angle of greater than or equal to 60° (0), less than or equal to 45° (1), less than or equal to 20° (2). (Jalil & Janvier, 2005:83; Tsuji, 2013:72; Turner *et al.,* 2015:92; Van den Brandt *et al.,* 2023:92)

93. Ectepicondyle, form: narrow and rounded (0); preaxially expanded wide rectangular flange (1). (Lee, 1997:82; Jalil & Janvier, 2005:84; Tsuji, 2013:73; Turner *et al.,* 2015:93)

94. Ectepicondylar foramen: absent (0); present (1). (Jalil & Janvier, 2005:85; Tsuji, 2013:74; Turner *et al.,* 2015:94; Van den Brandt *et al.,* 2023:94)

95. Entepicondyle, form: postaxially expanded wide rectangular flange (0); narrow and rounded (1). (Jalil & Janvier, 2005:86; Tsuji, 2013:75; Turner *et al.,* 2015:95; Van den Brandt *et al.,* 2023:95)

96. Entepicondylar foramen, form: completely enclosed (0); an ’open groove’ (1). (Jalil & Janvier, 2005:87; Tsuji, 2013:76; Turner *et al.,* 2015:96; Van den Brandt *et al.,* 2023:96)

97. Entepicondylar foramen, location: exposed in distal dorsal (extensor) view (0); ventral (flexor) view (1). (Jalil & Janvier, 2005:88; Tsuji, 2013:77; Turner *et al.,* 2015:97; Van den Brandt *et al.,* 2023:97).

98. Epicondylar distal projection: epicondyles do not project (far) past the radial and ulnar articulation surfaces (0); project past the radial and ulnar articulation surfaces, appearing ‘forked’ (1). (Jalil & Janvier, 2005:89; Tsuji, 2013:78; Turner *et al.,* 2015:98; Van den Brandt *et al.,* 2023:98)

99. Entepicondyle and ectepicondyle relative size: equal (0); ectepicondyle reduced (1). (Turner *et al.,* 2015:99; Van den Brandt *et al.,* 2023:99)

100. Radial condyle of the humerus, location: entirely ventral (0); more terminally, encroaches onto the distal end of the humerus (1). (Jalil & Janvier, 2005:92; Tsuji, 2013:81; Turner *et al.,* 2015:100; Van den Brandt *et al.,* 2023:100)

101. Humerus, ulnar articulation surface, form: groove bordered posteriorly by a faint ridge (0); groove bordered posteriorly by a prominent tubercle (1). (Jalil & Janvier, 2005:91; Tsuji, 2013:80; Turner *et al.,* 2015:101; Van den Brandt *et al.,* 2023:101)

102. Humerus, intercondylar depression, transverse ridge (intercondylar ridge border): on the distal dorsal end of the humerus, a transverse ridge separating the ulnar fossa (intercondylar depression) from the articulation (olecranon fossa) surface is absent (0); present (1). (Jalil & Janvier, 2005:90; Tsuji, 2013:79; Turner *et al.,* 2015:102; Van den Brandt *et al.,* 2022:102 modified; Van den Brandt *et al.,* 2023:102)

103. Humerus, ulnar fossa (intercondylar depression), depth: shallow depression (0); deep fossa (1). (Turner *et al.,* 2015:103; Van den Brandt *et al.,* 2023:103)

104. Humerus, ulnar fossa (intercondylar depression), width: ulnar fossa is much wider than the olecranon process (0); ulnar fossa is ‘narrow’, same width as olecranon process (1). (Turner *et al.,* 2015:104; Van den Brandt *et al.,* 2023:104)

**Ulna (characters 105, 106)**

105. Ulna, olecranon process, articulation surface: oriented medially (0); oriented terminally

(1). (Jalil & Janvier, 2005:93; Tsuji, 2013:82; Turner *et al.,* 2015:105; Van den Brandt *et al.,* 2023:105)

106. Ulna, olecranon process, size: well developed, greatly expanded past the most preaxial surface of the proximal articulation surface (0); reduced, nearly level with preaxial surface of the proximal articulation surface (1). (Jalil & Janvier, 2005:93; Tsuji, 2013:82; Turner *et al.,* 2015:106; Van den Brandt *et al.,* 2023:106)

**Manus (character 107)**

107. Manus, phalangeal formula: 23452, not reduced (0); 23332 reduced (1). (Jalil & Janvier, 2005:94; Tsuji, 2013:83; Turner *et al.,* 2015:107; Van den Brandt *et al.,* 2023:107)

**Sacral ribs (character 108)**

108. Sacral ribs, second and third: dorsoventral compression is slight (0); strong and sheet-like (1). (Jalil & Janvier, 2005:95; Tsuji, 2013:84; Turner *et al.,* 2015:108; Van den Brandt *et al.,* 2023:108)

**Pelvis (characters 109-119)**

109. Ilium, crista sacralis: weakly developed (0); well developed (1). (Jalil & Janvier, 2005:96; Tsuji, 2013:85; Turner *et al.,* 2015:109; Van den Brandt *et al.,* 2023:109)

110. Ilium, blade, expansion: not or only slightly anteriorly (0); well anteriorly (1). (Jalil & Janvier, 2005:98; Tsuji, 2013:87; Turner *et al.,* 2015:110; Van den Brandt *et al.,* 2023:110)

111. Ilium, anterior margin, lateral eversion: flat or slightly everted (0); surface strongly everted, oriented almost horizontal (1). (Jalil & Janvier, 2005:99; Tsuji, 2013:88; Turner *et al.,* 2015:111; Van den Brandt *et al.,* 2023:111)

112. Ilium, posterior process: long (0); strongly reduced (1). (Jalil & Janvier, 2005:100; Tsuji, 2013:89; Turner *et al.,* 2015:112; Van den Brandt *et al.,* 2023:112)

113. Ilium, shaft, orientation: vertical or posterodorsally inclined (0); anterodorsally inclined, forming an angle with the vertical of more than 20° (1); inclined even further anteriorly, forming an angle of more than 45° with the vertical (2). (Jalil & Janvier, 2005:97; Tsuji, 2013:86; Turner *et al.,* 2015:113; Van den Brandt *et al.,* 2023:113)

114. Acetabulum, dorsal buttress: not well developed (0); strongly developed (1). (Jalil & Janvier, 2005:101; Tsuji, 2013:90; Turner et *al.,* 2015:114; Van den Brandt *et al.,* 2023:114)

115. Acetabulum, anterior shape: round (0), notched (1). (Jalil & Janvier, 2005:102; Tsuji, 2013:91; Turner *et al.,* 2015:115; Van den Brandt *et al.,* 2023:115)

116. Pubis, process on the anterior border: absent (0); present (1). (Jalil & Janvier, 2005:105; Tsuji, 2013:93; Turner *et al.,* 2015:116; Van den Brandt *et al.,* 2023:116)

117. Pubis, median process: absent (0); present (1). (Jalil & Janvier, 2005:106; Tsuji, 2013:94; Turner *et al.,* 2015:117; Van den Brandt *et al.,* 2023:117)

118. Pelvic symphysis, length: long (0), short (1). (Jalil & Janvier, 2005:104; Tsuji, 2013:92; Turner *et al.,* 2015:118; Van den Brandt *et al.,* 2023:118)

119. Pelvic symphysis, dorsoventral thickness: thin (0), thick (1). (Jalil & Janvier, 2005:104; Tsuji, 2013:92; Turner *et al.,* 2015:119; Van den Brandt *et al.,* 2023:119)

**Femur (characters 120-125)**

120. Femur, head, preaxial expansion: no curvature (0); slight (1); strong (2). (Lee, 1997:107; Jalil & Janvier, 2005:107; Tsuji, 2013:95; Turner *et al.,* 2015:120)

121. Femur, trochanter major: absent (0); present (1). (Jalil & Janvier, 2005:109; Tsuji, 2013:96; Turner *et al.,* 2015:121; Van den Brandt *et al.,* 2023:121)

122. Femur, trochanter major, form: small, slightly thickened (0); large, more distinct (1). (Jalil & Janvier, 2005:109; Tsuji, 2013:96; Turner *et al.,* 2015:122; Van den Brandt *et al.,* 2023:122)

123. Femur, postaxial flange, length: limited to proximal region (0); extends entire length of femur (1). (Jalil & Janvier, 2005:112; Tsuji, 2013:97; Turner *et al.,* 2015:123; Van den Brandt *et al.,* 2023:123)

124. Femur, postaxial flange, width: narrows in the middle of the length of the femur (0); consistently wide, appearing straight (1). (Jalil & Janvier, 2005:112; Tsuji, 2013:97; Turner *et al.,* 2015:124; Van den Brandt *et al.,* 2023:124)

125. Femur, internal trochanter, shape: in ventral view, appears straight (0); proximally curved preaxially (1). (Jalil & Janvier, 2005:114; Tsuji, 2013:98; Turner *et al.,* 2015:125; Van den Brandt *et al.,* 2023:125)

**Tibia (character 126)**

126. Cnemial crest: cnemial crest of the tibia (longitudinal ridge on the dorsal (lateral or external) surface of the tibia) well developed and prominent (0); ridge and accompanying groove much reduced (1). (Jalil & Janvier, 2005:115; Tsuji, 2013:99; Turner *et al.,* 2015:126; Van den Brandt *et al.,* 2023:126)

**Tarsus (character 127)**

127. Astragalus and calcaneum: separate or sutured (0); fused, with the presence of the obturator foramen (1). (Jalil & Janvier, 2005:116; Tsuji, 2013:100; Turner *et al.,* 2015:127; Van den Brandt *et al.,* 2023:127)

**Pes (characters 128-131)**

128. Pes, phalangeal formula: 23454 or 23453 (0); 23343 (1). (Jalil & Janvier, 2005:118; Tsuji, 2013:101; Turner *et al.,* 2015:128; Van den Brandt *et al.,* 2023:128)

129. Pes, fifth digit: large, always longer than the first pedal digit (0); reduced, slender, shorter than the first pedal digit (1). (Jalil & Janvier, 2005:119; Tsuji, 2013:102; Turner *et al.,* 2015:129; Van den Brandt *et al.,* 2023:129)

130. Pes, metapodial (metacarpal and metatarsal), shape: slender, close to two times as long as wide (0); robust, approximately as wide as long (1). (Lee, 1997:120; Jalil & Janvier, 2005:120; Tsuji, 2013:103; Turner *et al.,* 2015:130)

131. Pes, non-terminal phalanges, shape: slender, 50% longer than wide (0); short, as long as wide (1); even shorter and more massive, about two times as wide as long (2). (Lee, 1995:46, Lee, 1997:121; Jalil & Janvier, 2005:121; Tsuji, 2013:104; Turner *et al.,*

2015:131)

**Osteoderms (characters 132-138)**

132. Osteoderms, body coverage: Osteoderms absent on the body (0); present (1). (Jalil & Janvier, 2005:122; Tsuji, 2013:105; Turner *et al.,* 2015:132; Van den Brandt *et al.,* 2023:132)

133. Osteoderms, body coverage, extent: osteoderms form only a longitudinal band closely overlying the vertebral column (0); cover entire dorsal surface of the body including flanks (1). (Jalil & Janvier, 2005:122; Tsuji, 2013:105; Turner *et al.,* 2015:133; Van den Brandt *et al.,* 2023:133)

134. Osteoderms, appendages: no osteoderms over the appendages (0); fore and hind limbs covered with numerous conical osteoderms (1). (Jalil & Janvier, 2005:127; Tsuji, 2013:110; Turner *et al.,* 2015:134; Van den Brandt *et al.,* 2023:134)

135. Osteoderm, appearance: dorsal surface of the osteoderms smooth, convex, without a central boss (0); possess a distinct rounded central boss (1); central boss on osteoderm capped by a small conical spine (2). (Jalil & Janvier, 2005:123; Tsuji, 2013:106; Turner *et al.,* 2015:135; Van den Brandt *et al.,* 2023:135)

136. Osteoderm, ornamentation: external surface of the osteoderms smooth and without ornamentation (0); osteoderms ornamented with fine, straight, regularly spaced ridges radiating out from a central boss to the edge (1); ridges fewer, larger, lumpier, and less regularly spaced (2). (Jalil & Janvier, 2005:124; Tsuji, 2013:107; Turner *et al.,* 2015:136; Van den Brandt *et al.,* 2023:136)

137. Osteoderm, dimension: round and small, with a dimension no larger than diameter of the centra of the dorsal vertebrae (0); osteoderms large, with a maximal length the same as or larger than the dorsal vertebral centra (1). (Jalil & Janvier, 2005:125; Tsuji, 2013:108; Turner *et al.,* 2015:137; Van den Brandt *et al.,* 2023:137)

138. Osteoderm, position: osteoderms do not touch, separated by a space (0); osteoderms more densely packed, often touching one another, but touching only on the shoulder and pelvic regions, never sutured or articulated over the trunk (1); osteoderms overlapping, articulated or sutured, forming a continuous layer on the dorsal surface of the body (2). (Jalil & Janvier, 2005:126; Tsuji, 2013:109; Turner *et al.,* 2015:138; Van den Brandt *et al.,* 2023:138)

**Gastralia (character 139)**

139. Gastralia: present (0); absent (1). (Jalil & Janvier, 2005:128; Tsuji, 2013:111; Turner *et al.,* 2015:139; Van den Brandt *et al.,* 2023:139)

**Extra (character 140-142)**

140. Marginal teeth, at mid region of maxilla or jaw, dorsoventrally tall (root-apical/mesiodistal ratio = 1.5 or higher), present (0), or absent (1). (Cisneros *et al.,* 2021:140; Van den Brandt *et al.,* 2023:140; Van den Brandt *et al.,* 2023:140)

141. Premaxillary tooth number, three (0), four or more (1), or two (2). (Cisneros *et al.,*

2021:141; Van den Brandt *et al.,* 2023:141; Van den Brandt *et al.,* 2023:141)

142. If the tabular (=supernumerary bone) is dorsally expanded, it does not contact the parietal (0), or it contacts the parietal (1). (Cisneros *et al.,* 2021:142; Van den Brandt *et al.,* 2023:142; Van den Brandt *et al.,* 2023:142)

---

**TABLE 1.** Measurements of the vertebrae of *Yinshanosaurus angustus* (IVPP V33181). Unit of measurement given in mm. *Abbreviations*: cen, centrum; ns, neural spine; pap, parapophysis; ver, vertebra

| NO. | height of ver | width of ver | length of ns | width of ns | height of ns | length of cen | width of cen | length of the pap |
| --- | --- | --- | --- | --- | --- | --- | --- | --- |
| cervical 3 | / | / | / | 43 | 82 | / | / | / |
| cervical 4 | >155 | 60 | 20 | >36.5 | >64.2 | 52 | 59 | ? |
| cervical 5 | >189 | 61 | 23 | >37.3 | >84.4 | 60 | 58 | ? |
| dorsal 1 | 211 | 80 | 26 | 53 | 94 | / | / | 23 |
| dorsal 2 | >205 | 97 | 34 | 57 | 77 | / | / | 38 |
| dorsal 3 | 200 | 190 | 35 | 55 | 78 | / | / | 41 |
| dorsal 4 | 210 | 180 | 35 | 52 | 82 | / | / | 37 |
| dorsal 5 | 218 | 170 | 42 | 51 | 70 | / | / | 42 |
| dorsal 6 | / | 195 | 44 | 40 | 70 | 49 | / | 36 |
| dorsal 7 | / | 193 | 48 | 41 | 67 | 50 | / | 39 |
| dorsal 8 | 275 | 160 | 42 | 34 | 73 | 60 | 60 | 28 |
| dorsal 9 | / | 160 | 72 | 33 | 70 | / | / | / |
| dorsal 10 | / | 162 | 77 | 32 | 79 | / | / | / |
| dorsal 11 | / | 155 | 80 | 29 | 76 | / | / | / |
| dorsal 12 | / | 156 | 91 | 28 | 87 | / | / | / |
| dorsal 13 | / | 158 | 88 | 28 | 82 | / | / | / |
| dorsal 14 | / | 160 | ? | 28 | / | / | / | / |
| dorsal 15 | / | / | / | 27 | / | / | / | / |
| dorsal 16 | / | / | / | / | / | / | / | / |

Unit of measurement given in mm

## Notes

### Competing Interest Statement

The authors have declared no competing interest.

